# Frequency modulated semi-continuous wave Doppler ultrasound: A novel method for the detection of scatterer velocity and range using wearable ultrasound instrumentation

**DOI:** 10.64898/2026.10.19.759949

**Authors:** Connor Delaney, Ahmed Bashatah, Abhishek Aher, Erica L King, Zahra Taghizadeh, Parag V Chitnis, Siddhartha Sikdar

## Abstract

Doppler ultrasound imaging has become a key tool for clinicians and researchers to measure the velocity of blood and tissues within the human body. Current clinical instruments utilize pulsed wave or continuous wave transmissions for tracking phase shifts or frequency shifts respectively to estimate scatterer motion. However, these two techniques can also be considered as ends of a continuum along which other motion tracking techniques exist. In this paper, we explore frequency modulated semi-continuous wave (FMSCW) Doppler, an underexplored Doppler imaging technique that has particular benefits for emerging wearable ultrasound applications due to significant simplification of instrumentation and lowered data rates. We present a full technical overview of this technique, simulations exploring the impact of selected imaging parameters as well as how noise impacts imaging, and present three exemplary imaging applications in human participants using FMSCW Doppler implemented on a wearable ultrasound system. The results of the simulations evaluating the impact of imaging parameters demonstrate the trade-off between resolution in the range and velocity axis as well as the impact of scatterer range and speed on the measurement error. The three sets of images collected from human participants serve to demonstrate FMSCW Doppler used in a variety of wearable ultrasound applications including cardiovascular monitoring, rehabilitation assessment, and prosthetic device control. Furthermore, we discuss the relationship between FMSCW Doppler and isomorphic techniques from radar as well as potential inspiration for improvements for FMSCW Doppler inspired by the echolocation of bats.

## I. Introduction

Since the first use of Doppler ultrasound for medical applications in 1957, distinct imaging paradigms have been developed to assess blood flow and the movement of different tissues for a wide range of different clinical applications [1]. Current clinical ultrasound systems implement different modes of visualizing flow and tissue motion, which fall into two broad categories: one utilizing short-duration pulsed wave (PW) transmissions and the second using continuous wave (CW) transmissions. PW builds upon the near ubiquitous pulse-echo imaging paradigm and tracks phase shifts in received pulses to determine velocity which typically is displayed as a spectrogram of flow velocity over time or mapped onto a color flow image. CW instead employs a separate transducer for transmission and reception and measures the frequency shift due to moving scatterers as a consequence of the Doppler principle. Note that in this paper, we will use the term “Doppler” to refer to velocity estimation and imaging following colloquial use in the field, although PW methods utilize phase shift estimation and not the frequency shift associated with the Doppler principle [2]. Both PW and CW techniques have benefits and drawbacks. PW can be used in conjunction with B-mode images of tissue structures while simultaneously measuring the velocity of these structures and flow inside blood vessels. PW methods often require more complex instrumentation and high sampling rates and are also limited by aliasing. CW Doppler on the other hand can be implemented with simple instrumentation, involves processing on demodulated audio-frequency signals, and theoretically is not affected by aliasing but loses the ability to resolve the depth, or range, at which scatterer velocities occur.

The problem of resolving the range and velocity of moving targets has been thoroughly investigated in other fields such as radar or sonar, and through natural evolution in the echolocation of bats and other animals [3], [4], [5], [6]. By drawing inspiration from these different fields, in this paper we investigate the premise that PW and CW approaches can be thought of as two ends of a continuum of range and velocity estimation approaches using ultrasound. Between PW and CW along this continuum other techniques with characteristics of both PW and CW can be designed by shaping the transmitted waveform and can be utilized to provide different tradeoffs between instrumentation and processing complexity and functionality. These approaches have particular relevance for motion estimation using wearable ultrasound instrumentation, an emerging paradigm shift in the field of ultrasound.

In this study we present an exploration of a method called frequency modulated semi-continuous wave (FMSCW) Doppler imaging, that exists between PW and CW Doppler along the continuum. This underexplored technique is both an extension of Time Delay Spectrometry (TDS) and an isomorphic method to frequency modulated continuous wave radar [7], [8]. TDS is an alternative imaging paradigm from the more conventional pulse-echo imaging which was first described by Richard Heyser in 1967 for acoustical measurements but later adapted to ultrasound imaging [7]. The foundational principle of TDS is that transmitted pulses consisting of a long (relative to that of pulse-echo pulses) linearly swept chirp signal and backscattered echoes shifted in frequency from the transmitted signal mix at the transducer.

This mixed signal can be demodulated with respect to the transmitted signal to recover an intermediate signal, called the beat signal, which encodes the depth of scatters in frequency content. This frequency content is then evaluated by simply applying the Fourier transform. Our research group has previously demonstrated the utilization of TDS ultrasound to measure muscle contractions with a multichannel wearable platform [9]. This system makes use of simplified hardware in comparison to conventional pulse-echo devices as the receive signal chain of a TDS imaging system is akin to a frequency-modulated radio receiver and the baseband signal requires a much lower rate of digitization than conventional pulse echo systems (typically kHz rather than MHz). These differences lead to lowered memory and power requirements further lowering the hardware requirements of the system. These benefits directly aid in the packaging of ultrasound imaging systems into wearable platforms and are made possible due to the TDS imaging technique. However, by taking inspiration from isomorphic techniques in the field of radar this system and others using the same imaging paradigm can also perform Doppler imaging without modifying any of their hardware.

The isomorphism between radar (as well as sonar) and Doppler ultrasound imaging is well established due to them operating along similar physical principles [8], [10]. Effectively both fields employ methods for the estimation of object distance and velocity based on transmission and reception of electromagnetic or pressure waves. This makes processing techniques used in one field often directly translatable to the other. With respect to the present work, the radar technique which inspires FMSCW Doppler is frequency modulated continuous wave (FMCW) radar [4]. This technique utilizes linearly swept chirp transmissions across a specified bandwidth and then repeats periodically creating a sawtooth pattern or triangular pattern if alternating up and down chirps are utilized. Using a homodyne reception scheme, the transmit and backscattered receive signals are mixed and then lowpass filtered to produce a beat signal or alternatively referred to as the intermediate frequency signal. This beat signal encodes changes in range and velocity of scatterers as changes in frequency and phase respectively [4].

FMCW methods have been applied to ultrasound by a few research groups but remain widely underexplored. Initial investigations into this Doppler technique using frequency modulation were performed by Wilhjelm and Pedersen who probed the theory underlying the technique and performed experiments and simulations to determine how this new technique compared to PW Doppler [11], [12]. These initial studies observed similar performance between PW and their implemented frequency modulated Doppler. Following this, the potential of FMCW Doppler radar principles being coopted for use with ultrasound systems was expressed by Kunita *et al*. [13]. This team extended the work of Wilhjelm and Pedersen with improvements to the demodulation process used in gathering the beat signal as well as performing evaluations with experiments imaging phantoms. Furthermore, this team also probed the possibility of measuring the range of scatterers with FMSCW Doppler ultrasound [14].

In this study, we present a thorough overview of estimating scatterer range and velocity using FMSCW Doppler followed by an evaluation of these techniques using simulations. We then demonstrate the implementation of this technique to record human tissue movement through several *in vivo* experiments. We conclude with a discussion of the FMSCW Doppler method, its limitations, and the exciting prospects of future developments based on the foundation presented in this study.

## II. Overview of Range and Velocity Estimation

This section details the mathematics used by FMSCW Doppler to take backscattered long-duration; frequency sweeping pulses and calculate range-resolved tissue velocities. Several approaches may be utilized to estimate the velocity of tissues from FMSCW Doppler, which are each presented in this section. The starting point for the derivation of FMSCW Doppler is a linearly swept frequency modulated chirp signal.

Consider a frequency modulated signal transmitted by the ultrasound transducer:

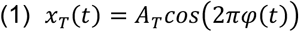

where *A*_*T*_ is the transmitted amplitude and *φ*(*t*) is the instantaneous phase. The instantaneous frequency *f*_*T*_ (*t*) is given by the derivative of instantaneous phase:

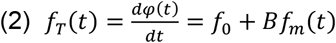

where *f*_0_ is the initial frequency and *f*_*m*_(*t*) is a slow-varying modulation function. *B* is a selected, adjustable factor that describes the degree of modulation. When *B* =0, there is no frequency modulation.

The modulation function *f*_*m*_(*t*) can be designed to have different properties. For example, the modulation function can be a sawtooth wave, with linearly increasing frequency during a period *T*_*s*_,

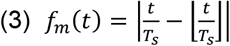

where 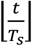 represents the floor function.

Or the modulation function *f*_*m*_(*t*) can have the form of a triangular wave, where the frequency increases linearly and decreases linearly in a periodic manner.

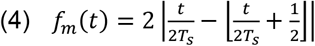

The instantaneous frequency of this modulation function is a linear chirp increasing from an initial frequency of *f*_0_ to a frequency of *f*_*e*_ (i.e., the up-chirp) and then decreasing linearly from *f*_*e*_ back to *f*_0_ (i.e., the down-chirp). In this case, the bandwidth, *B* = *f*_*e*_ − *f*_0_, is modulation factor *B*, and *T*_*s*_ is the duration of each up- and down-chirp sweep (i.e., sweep time). The ratio 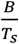 is then the sweep rate or the rate of frequency change in the transmitted signal. The bandwidth, sweep time, and sweep rate of the sawtooth modulation function can be found in the same manner, but rather than exhibiting both up- and down-chirps it consists of exclusively up-chirp sweeps.

As the acoustic wave travels through the medium and interacts with a scatterer at a distance *d* from the transducer moving at a constant velocity *v* in the imaging plane (i.e., towards or away from the transducer), the received backscattered signal with amplitude *A*_*R*_ is delayed in time, and the instantaneous frequency is Doppler shifted from the perspective of the transducer receiving the backscattered signal as shown in Fig 3. Thus, we can represent the received signal as:

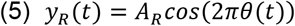

where:

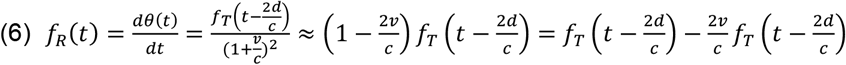

The transmitted and received signals can then be mixed to yield:

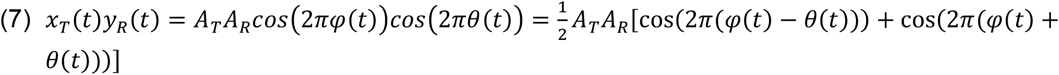

A low pass filter is then used to isolate the difference frequency and suppress the higher sum frequency resulting in an intermediate frequency signal:

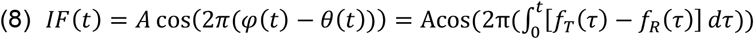

Substituting from Eq. 6, we get

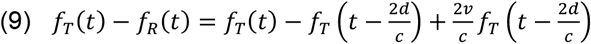

Substituting from Eq. 2, we get

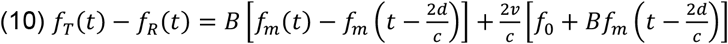

Using the case of a sawtooth modulation function in Eq. 3, consisting of an up-chirp repeating periodically, consider the up-chirp spanning from time interval [0, *T*_*s*_]. For the up-chirp in this interval,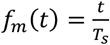 . Substituting into Eq. 8 and Eq. 10 yields

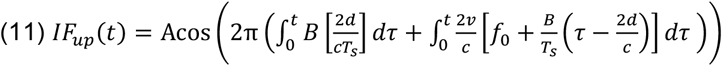

If we assume *d* and *v* to be constants, after performing the integration, this simplifies to:

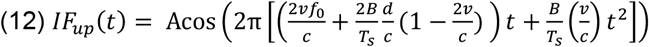

The demodulated intermediate frequency signals contain the range frequency,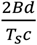 , and doppler shift frequencies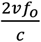, and the frequency varies with a sweep rate of 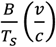.

It can be seen from Eq. 12 that if the scatterer velocity *v* is zero, then the intermediate frequency signal exclusively resolves range frequency shift 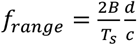 corresponding to the range of the scatterer *d*.

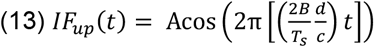

Conversely, if there is no frequency modulation (i.e., if *B* = 0) then the intermediate frequency signal exclusively resolves a Doppler frequency shift 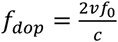 corresponding to the velocity of the scatterer *v*. This is identical to the behavior observed with CW.

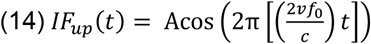

For a triangular modulation function (Eq. 4) with periodically repeating pairs of up- and down-chirps, the expression for the intermediate frequency resulting from the up-chirp remains the same. The corresponding downchirp in the interval [0, *T*_*s*_], can be obtained by substituting *f*

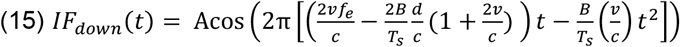

The range frequency shift has a negative sign relative to the up-chirp in Eq. 13, while the Doppler frequency shift has a positive sign. Fig 4 shows a graphical visualization of the transmitted and received instantaneous frequencies.

Choice of different sweep rates can further separate the range frequency shift and the Doppler frequency shift from one another. Range resolution depends on the sweep rate 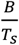 , with increasing sweep rates providing better frequency separation between two interfaces separated by a fixed distance. The Doppler resolution depends on the initial chirp frequency and the pulse duration *T*_*s*_. Specifically, the ability to resolve frequency shifts is related to the time-bandwidth product. Longer duration pulses have better frequency resolution both for range and Doppler shift.

### Signal Analysis Methods

Digital signal processing can be used to distinguish the range and Doppler shift frequencies from one another in order to resolve them independently. Different methods for resolving range and Doppler shift are relevant for slow and fast movements.

Motion in the human body happens at much smaller speed (e.g., blood flow is less than 1 m/s and tissue motion is frequently less than 10 cm/s) compared to the speed of sound in tissue (∼1540 m/s).

Thus, for a chirp with starting frequency of 3MHz, the Doppler shift frequency,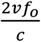 , is less than 4kHz for blood flow and less than 400 Hz for tissue motion. In contrast the frequency shift corresponding to the range,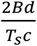 , depends upon the choice of sweep rate and the range of the scatterer of interest. For example, using a sweep with 3-5MHz bandwidth=*B* over 10ms, the range frequency shift for a scatterer at 5 cm depth is ∼13 kHz. Therefore, for tissue motion the range frequency shift is often an order of magnitude higher than the Doppler frequency shift. Analysis to resolve range and velocity of scatterers can be performed in the frequency domain or the time domain using different approaches, each with their own practical applications.

#### Frequency domain analysis using the 2D Power Spectral Density of the Intermediate Frequency Signal

Consider a transmitted ultrasound signal is frequency modulated with a triangular waveform such that the transmitted signal consists of a linear up-chirp from frequency *f*_*0*_ to *f*_*e*_ over *T*_*s*_ seconds, and a linear down chirp from frequency *f*_*e*_ to *f*_*0*_ over *T*_*s*_ seconds. The received demodulated intermediate frequency signal *IF*(*t*) is digitized at a rate *F*_*r*_ (i.e., a sample is obtained every 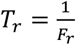 seconds). Now consider a scatterer at depth *d*, moving at a velocity *v*. In the human body, the scatterer movement is expected to be time varying, so both the instantaneous depth and velocity are functions of time. Thus, from Eq. 11, for the up-chirp we obtain

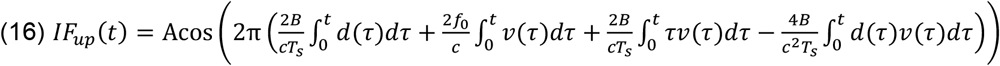

The factor 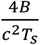 is very small relative to the other terms, so the expression can be simplified to

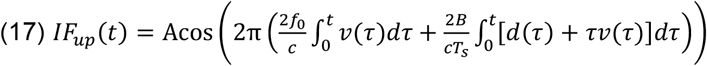

A similar expression can be derived for the down-chirp.

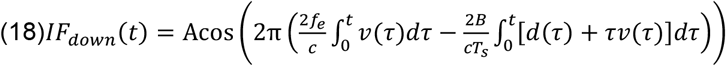

We can reorganize the digitized signal into a 2D signal consisting of set of “fast time” samples obtained at times, *nT*_*r*_, that denotes time within the duration of a single given chirp, and a set of “slow time” samples obtained at times, *kT*_*s*_, that denotes the samples recorded at the same time within chirps across multiple received chirp signals.

Although the expressions in Eq. 17 and 18 represent the general case, for illustrative purposes we can make a simplifying assumption that the scatterer position *d(t)* and velocity *v(t)* are slowly varying relative to the duration of the chirp *T*_*s*_. Thus, in fast time, the position and velocity are considered to be constant *d(kT*_*s*_) and *v(kT*_*s*_) within the duration of the chirp. We then obtain

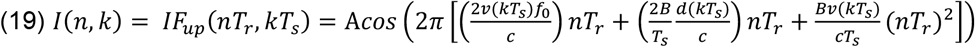

The 2D power spectral density of a windowed portion of this demodulated intermediate frequency signal can be computed using the 2D Fourier Transform

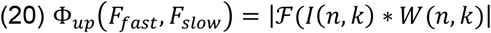

Where *W* is a 2D window function used to select the portion of the intermediate frequency signal for analysis. The characteristics of *W* also impact the range and Doppler resolution, since its qualities are reflected in the 2D power spectral density.

The spectral centroids identify the center of energy within the 2D spectrum providing a measure of the aggregate focal point of activity in both range and velocity. Spectral centroids of the 2D power spectral density are given by

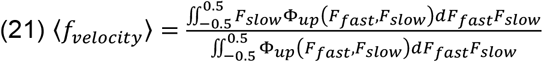

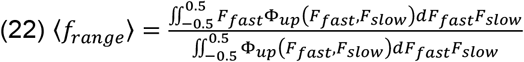

As with other Doppler imaging techniques, clutter filters and depth gates may be applied before the calculation of the spectral centroid as befitting a given application to focus measurements on specific regions and velocities. Furthermore, the 2D power spectral density can be projected on to either the range or velocity axes, a view of the data which may be well suited for some applications.

#### Disambiguation of Velocity Centroid to reduce impact of Aliasing

Aliasing is a limitation that plagues PW and drives increasing hardware requirements for imaging systems in order to combat it. When the frequency content in the velocity axis associated with a scatterer exceeds resolvable frequency based on the pulse repetition rate, the signal corresponding to that scatterer becomes aliased and appears on to the opposite side of the spectra. One way of visualizing this is the frequency spectra wrapping around and meeting at its extremes such that a positive velocity exceeding the maximum resolvable velocity appears as a negative velocity. In principle the velocity can wrap around range of resolvable frequencies multiple times, which make counteracting aliasing more difficult in high acceleration situations. However, in situations with velocities that do not alias beyond the zero of the velocity axis, unwrapping and disambiguating the velocity signal is relatively straightforward.

Using the example of a velocity spectrogram where the information from the 2D Fourier transform has been collapsed onto the velocity axis, divide the frequency bins of the Doppler axis into four segments of equal length. Then concatenate a copy of the spectra within the negative most segment of the velocity axis onto the positive most region. Perform the same operation by copying the positive most segment of the spectra and concatenating it onto the most negative region of the velocity spectra. This operation has effectively “unwrapped” the velocity spectrogram such that one is able to observe velocity information that would alias in the proper position along the velocity axis.

The disambiguation operation requires one to find the centroid in the range dimension as described previously. Taking the derivative of this range centroid provides an approximation of the velocity signal. This can be utilized to determine the direction of the velocity signal. Comparing the velocity approximated from the range centroid to the noise floor, defined as twice the standard deviation of the five previous samples in the approximated velocity, at each time point of the spectrogram will determine which region of the unwrapped velocity spectrogram to zero thus removing it from the calculation of the velocity centroid. If the approximated velocity from the range centroid is greater than the noise floor for a given time point, set all frequency bins in the negative most third of the velocity axis to zero. If the approximated velocity is less than the noise floor, set all frequency bins in the positive most third of the velocity axis to zero. Then the velocity centroid can be calculated as described previously using the velocity spectrogram with regions with potential aliasing removed.

The same premise may be applied to 2D spectra used to calculate the centroid. Simply unwrap each 2D spectra in the velocity dimension and then use the range centroid to zero the third of the unwrapped velocity axis which is not to be used for the velocity centroid calculation.

#### Imaging Velocities at all Ranges using the Color M-Mode

Just as it is possible to project the 2D power spectral density onto either the range or velocity axes, it is also possible to generate a M-mode image that presents the net velocity at each depth rather than the intensity of the backscattered signal at each depth. This can be calculated for each frame using

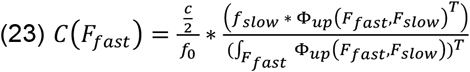

Where *f*_*slow*_ is the vector of slow time frequencies resolved by the Fourier transform and *T* is a transpose operation. When this calculation is performed for all frames, a color M-mode is produced displaying the net velocities at all imaged depths across time. This view of scatterer velocities may improve interpretability when multiple scatterers are present as well as providing another view to inform where range gates should be applied.

#### Relationship of FMSCW to PW and CW

The relationship between FMSCW Doppler and conventional PW and CW methods is perhaps best conceptualized in the frequency domain considering a 2D Fourier transform along both fast-time (i.e., within the pulse) and slow-time (i.e., inter-pulse) dimensions, as shown in Figure 1 [4], [15], [16]. In the case of FMSCW Doppler ultrasound, the 2D Fourier transform approach estimates the range and velocity of scatterers by extracting the frequency and phase shifts respectively due to each scatterer. The contributions of frequency and phase shifts due to scatterer range and velocity are separable due to them differing by an order of magnitude and are measured by considering individual periods of transmitted sweeps as columns of the input to the 2D Fourier transform arranged in the order in which they were collected. The relationship between the 2D Fourier transform range-velocity estimator and TDS imaging becomes apparent when it is considered that the projection of the 2D Fourier transform onto the range dimension is identical to the A line produced from conventional TDS imaging. Kunita *et al*. used a similar principle by estimating the frequency content of the beat signal to determine the power at a given range [14]. For a detailed discussion on TDS ultrasound refer to [9] and for a description of FMCW radar refer to [4]. Note that our FMSCW method does not require a continuous transmission. We can estimate velocity from purely the up chirp or down chirp signals, and turn off transmission until the next pulse repetition interval, thus our approach is distinct from the velocity estimation approach of Kunita *et al*. [13].

**Figure 1.**
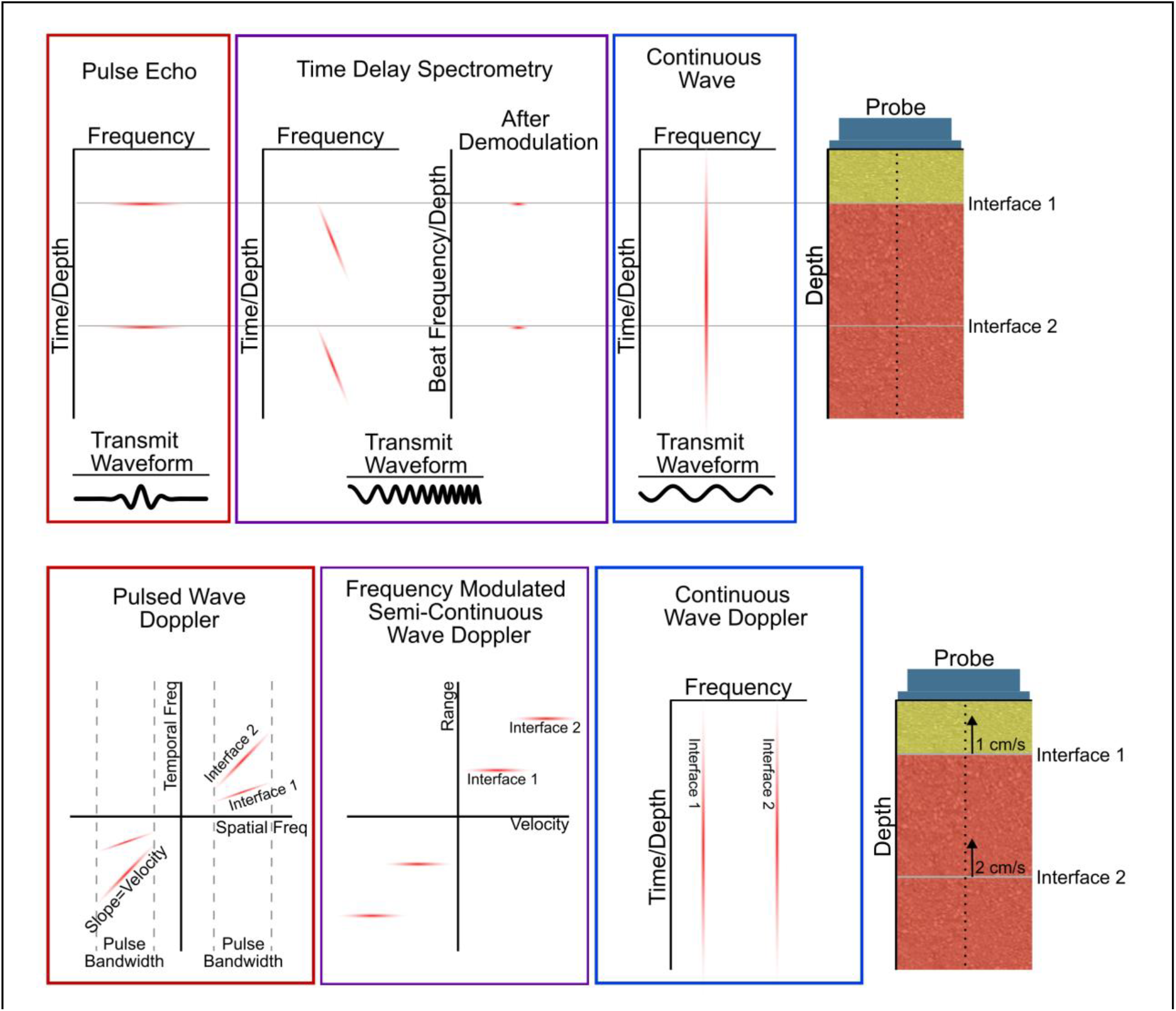
Illustration of the differences between PW, CW, and FMSCW Doppler and their associated standard imaging technique should they have one. **Top:** Illustration depicts a single element probe placed above a muscle collecting A-mode images. The yellow region represents the subcutaneous layer and the red region below the probe represents muscle tissue. One interface lies at the boundary of the subcutaneous layer and muscle, while the other is a fascia within the tissue. Each Doppler imaging technique has the time-frequency behavior of receive signals depicted including the FMSCW Doppler signal after demodulation. Note that CW cannot differentiate the depth of the two interfaces. Furthermore, the shape of the transmit signals for each technique is presented below the illustration of the receive signals. **Bottom:** Illustration depicts a single element probe performing Doppler imaging. In this illustration each of the two interfaces have different velocities moving towards the probe. Illustrated plots present the behavior observed in the 2D Fourier transform for PW and FMSCW Doppler and time-frequency behavior for CW Doppler.

**Figure 2.**
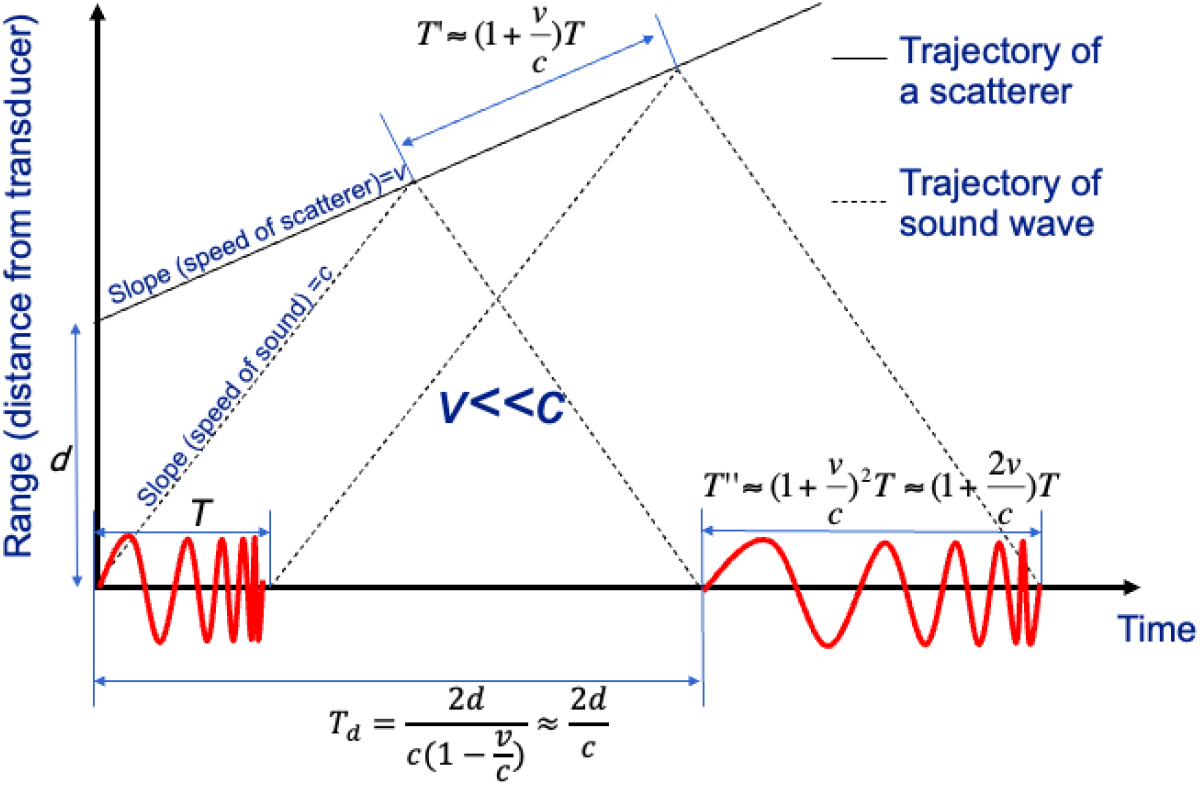
Visual representation of a frequency modulated signal transmitted by an ultrasound transducer at time t=0 interacting with a scatterer at an initial depth of d. The ultrasound signal travels at the speed of sound, c, while the scatter is traveling at a speed, v away from the transducer. As a result of the interaction, the backscattered reflected signal is shifted in time proportionally to the received signal, and the instantaneous frequency is Doppler shifted proportionally to the travel speed.

**Figure 3.**
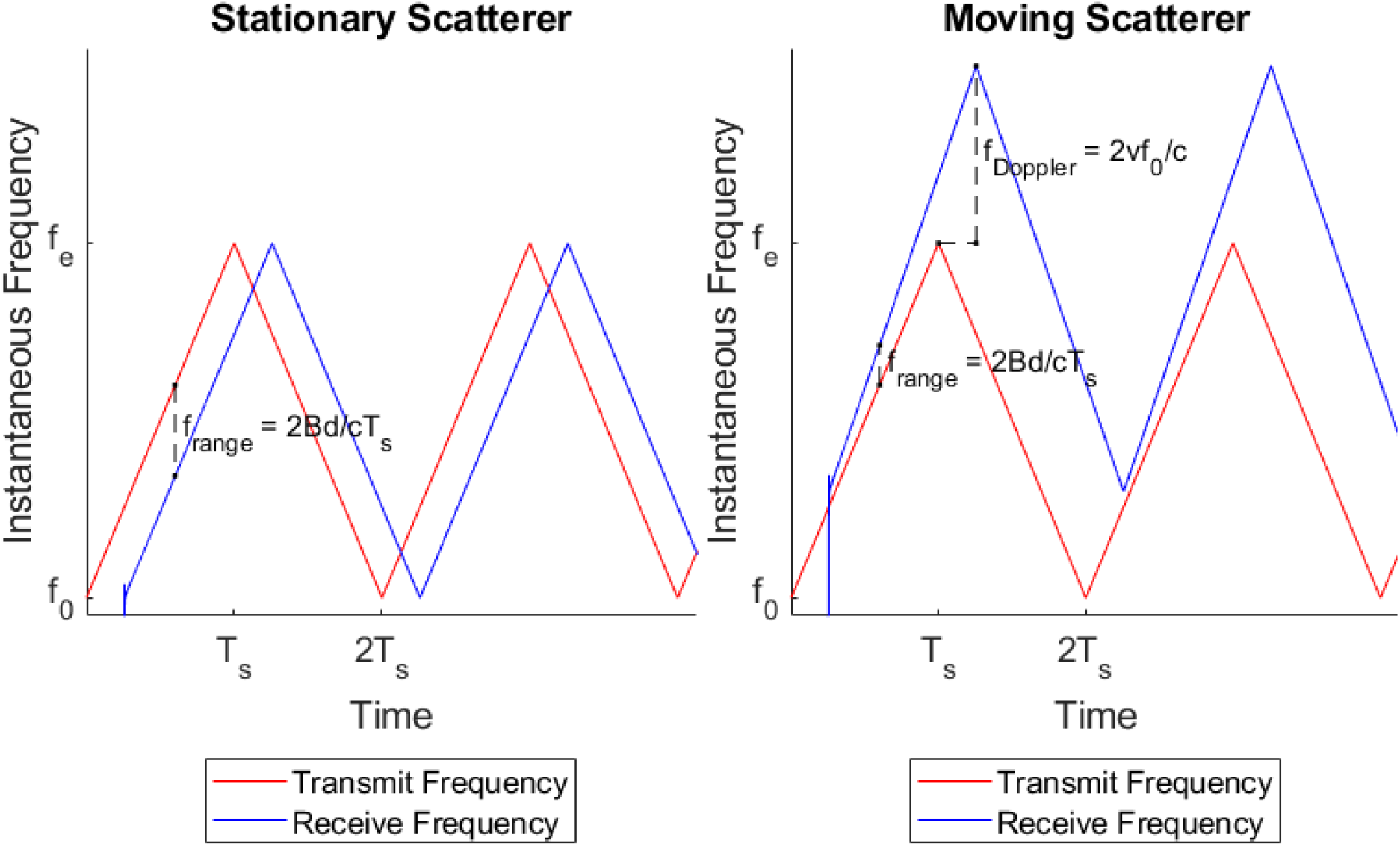
**Left:** The instantaneous transmit *f*_*T*_(*t*) and receive frequencies *f*_*R*_(*t*) for a triangular chirp signal (i.e., linear up- and down-chirp) in the case of a stationary scatterer at a depth *d*. The received signal is shifted in time, which corresponds to a frequency shift *f*_*range*_. **Right:** The instantaneous transmit and receive frequencies for a triangular chirp signal in the case of a scatterer at a depth *d* moving with a speed *v*. The received signal is shifted in time as well as Doppler shifted with frequency shift *f*_*dop*_. The Doppler shift and range shifts have been exaggerated for illustration

**Figure 4.**
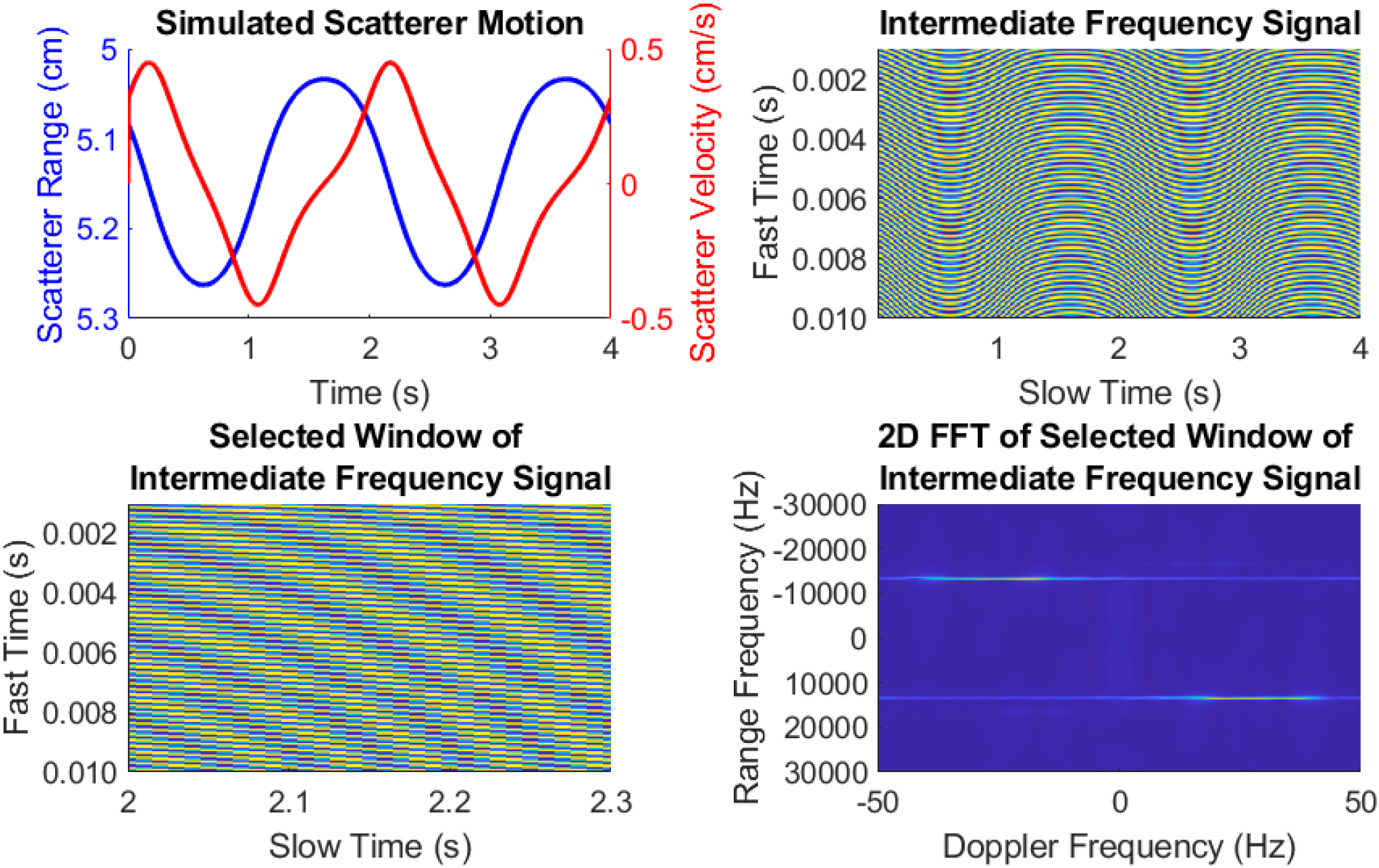
Simulation demonstrating the signal processing algorithms for estimating time varying scatter motion. **Top left**: A simulated scatterer at an initial depth of 50.5 mm, moving periodically every two seconds up to a depth of 52.7 mm. The maximum scatterer velocity is +/-0.45 cm/s. **Top right**: The scatterer is imaged using a triangular chirp signal with f_0_=3MHz, B=2MHz, and T_s_=10ms. The intermediate frequency signal can be represented as a 2D matrix, where the values in the rows of each column correspond to the samples of the demodulated signal within each chirp (fast time), and each column corresponds to consecutive chirps (slow time). **Bottom left**. A selected fast and slow time window of the intermediate frequency signal shows the variation in slow time corresponding to the maximum positive velocity of the scatterer. **Bottom right**. The corresponding 2D Fourier Transform of the 2D signal in the bottom left panel shows the Doppler frequency axis (corresponding to slow time) and the Range frequency axis (corresponding to fast time). In this case the range shift frequency is ∼13.3kHz, and the Doppler shift frequency is ∼30 Hz. Note that due to time-varying velocity within the analysis window, there is a spreading of the Doppler spectrum. The Doppler spectrum can be estimated by taking the projection along all range frequencies, while the range spectrum can be estimated by taking the projection along all the Doppler frequencies. Frequency domain filters can be applied before taking the projection to select specific range and Doppler frequency shifts of interest. The mean frequency shift (i.e., the tissue or blood flow velocity) can be estimated from the spectral centroid of the Doppler spectrum.

#### Time domain analysis using the 2D Autocorrelation of the Intermediate Frequency Signal

Computations in the time domain can also be used to extract the range and velocity of scatterers using the 2D or 1D autocorrelation applied to the reorganized digitized intermediate frequency signal in a manner not wholly dissimilar to other Doppler velocity estimators [17]. Consider the analytical signal 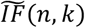 corresponding to the intermediate frequency signal *IF*(*n, k*) obtained using the Hilbert transform along the fast time direction. The 2D autocorrelation 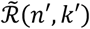 is then defined by

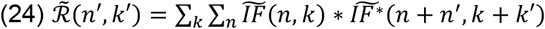

The 2D autocorrelation function 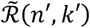 is also the inverse Fourier transform of the 2D power spectrum. The discrete derivatives of the 2D autocorrelation function can be shown to correspond to the spectral centroids of the 2D power spectrum, i.e.,

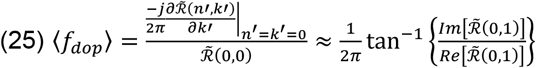

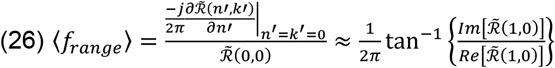

The estimated velocity can then be derived using

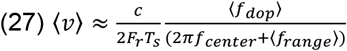

Where *f*_*center*_ is the center frequency of the chirp normalized by sampling rate of the intermediate frequency signal *F*_*r*_.

The principles of the 2D autocorrelation may also be applied to generate a 1D autocorrelation. Again this technique applies the premise utilized by other 1D autocorrelation velocity estimators [17]. In effect the 1D autocorrelator estimates the mean velocity using the velocity and the centroid of the analytic range spectrum at *f*_*center*_. The 1D autocorrelation is calculated as

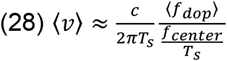

Both 1D and 2D autocorrelation techniques may be improved by incorporating a range gate to limit the scatterers which contribute to the estimated velocity. This can be readily performed by filtering out frequency content from ranges outside of the range gate. This filtering can either be performed in the frequency domain or the time domain. We demonstrate both methods in our results in the following sections.

Filtering in the time domain to apply a depth gate on the autocorrelation is straightforward as the frequency content in the beat signal is directly mapping the range and velocity of the scatterer. To determine the passband for a linear modulation function, determine the minimum and maximum depths and then use:

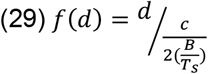

Where *d* is either the minimum or maximum depth for the depth gate. Any bandpass filter may be utilized in order to apply the depth gate, but for this study we opted to use a 4^th^ order Butterworth filter.

Filtering in the frequency domain can be performed during the process of calculating the analytical signal. By taking a Fourier transform of the beat signal for each frame, zeroing all frequencies outside the depths of interest, and then taking the inverse Fourier transform one is able to effectively apply a depth gate in the frequency domain. These filtered signals then may be used directly with Eq. 25 and 26.

#### Methods for Analysis of Fast-moving Blood Flow

For fast moving blood, where the velocity can be as high as 1m/s, we can no longer make the assumption that the velocity is constant within a pulse repetition interval. Therefore, we need to utilize the more general equation Eq. 17. However, the depth of the blood vessel can be assumed to be constant, so we obtain:

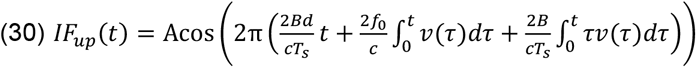

The instantaneous frequency of the demodulated up-chirp and down-chirp signals are therefore

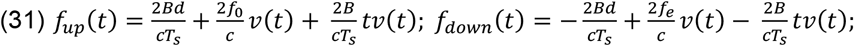

The instantaneous frequency can be estimated using spectral (frequency domain) methods, as well as autocorrelation (time domain) methods.

#### Waveform shaping of the transmitted chirp to improve estimation performance

The above method imposes fundamental tradeoffs between range and velocity resolution that can be improved by borrowing inspiration from nature. Bats utilize chirp signals for echolocation, but they do not use linear frequency modulated chirps at all times [3], [18]. Rather, they adjust the sweep rate nonlinearly to optimize range and velocity resolution. This adjustment is made based on proximity to prey, suggesting that predation needs are used to tune the chirp behavior. A similar principle can be applied to our theoretical framework as well.

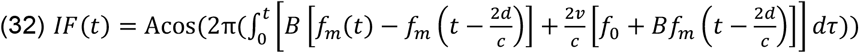

An example choice, inspired by bat echolocation may be a downchirp with 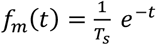

In this case, the intermediate frequency signal simplifies to

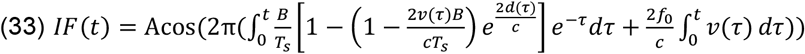

The parameters *B* and *T*_*s*_ can be chosen such the Doppler frequency shifts and range frequency shifts are significantly separated. The analysis can be performed by selecting different time windows within the period of the modulation function to isolate Doppler shift or range shift. Isolating Doppler and range shift further may provide improved range and velocity estimates. However, mimicking the real-time adjustments to chirp shape and bandwidth of bats is likely even more beneficial than simply selecting the same sweep behavior.

## III.Experiments

To evaluate the characteristics and performance of FMSCW Doppler imaging, we performed simulations probing the impact of selected linear sweep parameters and response to noise. Furthermore, we then utilized a wearable ultrasound imaging system to perform *in vivo* imaging of human tissue to demonstrate FMSCW applications. All human studies were performed following procedures approved by our Institutional Review Board.

### Simulations

#### Investigation of linear sweep parameters

The first simulation was conducted to characterize how linear sweep parameter selection impacts resulting images. All combinations of four sweep bandwidths and four sweep durations were simulated using the same motion for a resulting sixteen different sweep rates. The sweep bandwidths included .5MHz, 1MHz, 2MHz, and 4MHz while the sweep durations which were used included 1ms, 2.5ms, 5ms, and 10ms. The center frequency of the simulated sweeps was fixed at 4MHz. A fixed sampling rate of pulses as assigned as 400kHz for all permutations of the simulation. Linearly swept up-chirps (i.e., chirps with increasing frequency) were used for all transmitted signals to create a sawtooth pattern as in Eqn 3.

Receive signals were similarly simulated using delay and Doppler shift based on the movement of a single scatterer. This scatterer approximated the movement of muscle fascia during a small amplitude contraction and was utilized to evaluate the resulting resolution and accuracy of the velocity and range estimations. This scatterer was initially positioned at 2cm deep into the imaging plane with and moved towards the transducer with a maximum velocity of 3mm per second, held position, and then moved away from the transducer at the same rate to return to its initial position. The receive signals also had a 1.059MHz/cm (0.5dB/MHz/cm) attenuation coefficient applied to mimic tissue attenuation. Transmit and receive signals were mixed and lowpass filtered before being processed with FMSCW Doppler. For this simulation, the 2D Fourier transform was used to estimate the range and velocity of the scatterer for simulations of all combinations of sweep parameters. The full-width half-maximum (FWHM) of the 2D Fourier transform used to estimate velocity and range was calculated to determine the impact that sweep parameters have on the resulting resolution of the velocity and range estimation.

#### Investigation of the impact of noise

A second and third simulation were conducted to characterize how noise impacts characteristics of resulting images and the estimated velocity and range of scatterers. Specifically, the second simulation was aimed at determining the impact of noise as it relates to the selected sweep parameters, while the third simulation probed the impact of noise as it relates to the position and velocity of scatterers.

In the simulation investigating the relationship between noise and the selected sweep parameters of transmitted chirps, the selection of sweep parameters was the same as described in the section above. The position and movement of the single scatterer as well as the simulation of transmit and receive signals were performed in the same manner as the previous simulation as well. However, in this simulation Gaussian noise was added to the receive signal in the time domain with a power determined to provide a set signal-to-noise ratio (SNR). SNRs of -30, 0, 30dB, and no noise were used and applied to each combination of sweep parameters. The root mean square error (RMSE) was calculated for various velocity and range estimators for all combinations of sweep parameters at all SNRs.

For the third simulation targeting the relationship between scatterer movement and noise, rather than varying sweep parameters, the initial position and maximum velocity of scatterers was varied. Three initial ranges were used for the scatterer (15mm, 35mm, 65mm) and three maximum velocities were used (0.5cm/s, 1cm/s, 1.5cm/s) to create nine different combinations of scatterer motion and starting position. These combinations were simulated with Gaussian noise added to the receive signal in the same manner as described for the previous simulation at SNRs ranging from -30dB to 30dB. The RMSE for these simulations was calculated to evaluate the impact on estimators due to noise as it relates to the initial position and velocity of scatterers.

In addition to the RMSE for each velocity estimation method, the resolution of the 2D Fourier transform in both the range and velocity dimensions was calculated for all combinations of SNR and sweep parameters or scatterer motion for both simulations.

### Human Imaging

#### Hardware and software implementation of FMSCW Doppler with a wearable imaging system

To demonstrate the practical applications of FMSCW Doppler, we performed experiments on human subjects to measure the velocities of moving tissues *in vivo*. Ongoing work is exploring each of these applications, and in-depth studies of these applications with larger sample sizes are outside the scope of this paper. Building off of the previous work by our research group, a prototype system was developed to perform FMSCW Doppler imaging [9]. The prototype has four ultrasound single element transducers designed for generating M-mode images using a compact electronic driver/receiver circuit. Each ultrasound transducer is comprised of 1-3 piezocomposite lead zirconate titanate (PZT) and two copper-tin electrodes designed with a center frequency of 4MHz, acoustic impedance of 17 MRayl, and 7mm diameter. A polyvinylidene fluoride (PVDF; 110 µm thick, 4.2 MRayl impedance) matching layer and room-temperature-vulcanizing silicone backing layer was used to further improve transducer performance. The waveform produced by the driver circuit is generated by a programmable signal generator. For our purposes, a triangular waveform with linear up-chirp and down-chirp signal was implemented controlled via a microcontroller. Using faster sweep rate by decreasing the chirp duration will increase the range of detectable Doppler velocities without aliasing, with the trade-off of shorter chirp duration resulting in coarser range resolution (i.e., fewer samples in the fast time dimension of the intermediate frequency signal) and a lower SNR. The driver circuit produced a linear FM sweep from 3.6MHs to 4.3MHz over 2.5ms, corresponding to *f*_*0*_, *f*_*e*_, and *T*_*s*_ respectively.

The FMSCW Doppler prototype transmits the driving frequency modulated chirp signals simultaneously on all ultrasound transducers (referred to as channels). The demodulated intermediate frequency signals for all channels, consisting of frequencies in the audio frequency range, are lowpass filtered and digitized with a sampling frequency using the onboard analog to digital converter of a microprocessor (60kHz sampling rate, at 16 bits/sample). The digitized intermediate frequency signal can be processed locally on the microprocessor or transmitted via a communication interface to a central processing unit of a computer. For the purposes of the demonstration applications in this study, the intermediate frequency signal was transferred to a computer for offline processing.

#### Imaging human carotid wall movement

An emerging application of wearable ultrasound is the continuous estimation of blood pressure using the arterial pulse wave velocity and arterial wall distension [19], [20]. Thus, our first demonstration application is to evaluate the ability of our proposed method to measure carotid arterial wall movement. A single transducer from the FMSCW Doppler prototype imaging system described in the previous section was placed above the carotid artery of an individual without any known cardiovascular issues. Imaging of the movement of the arterial wall was performed at rest for 25 seconds. Both conventional M-mode images using TDS and FMSCW Doppler estimates of velocity using a 2D Fourier transform method were collected.

#### Imaging vastus lateralis movement during body weight squats

Another emerging application of wearable ultrasound is the dynamic monitoring of musculoskeletal function [21], [22]. Our second demonstration application of FMSCW Doppler consisted of measuring the movement of muscle fascia during a body weight squat in an individual one year after an anterior cruciate ligament (ACL) rupture and repair on their left side. Two transducers were placed on the vastus lateralis bilaterally while a participant performed ten body weight squats. As with the previous human imaging, both conventional M-mode images using TDS and FMSCW Doppler estimates of velocity and range were collected. The range and velocity were estimated using the 2D Fourier transform method with a range-gate around the primary fascia within the image. The resulting velocity traces were used to extract the average peak velocity for each side, and the cross-correlation of these two velocity traces was taken to find the time lag between peak contractions on each side and the symmetry of contractions at this maximal alignment. Furthermore, a color M-mode was produced using the 2D Fourier Transform.

#### Imaging attempted hand gestures in residual limb

Wearable ultrasound has been proposed for sonomyography, a method to recognize volitional gestures for the purpose of controlling human computer interfaces or an assistive device such as a prosthetic hand [23], [24]. Thus, a third demonstration application involved imaging the residual limb of an individual with trans-radial limb loss. For this demonstration, all four transducers of the FMSCW Doppler prototype were placed on the residual limb of the participant emulating a configuration for sonomyographic control of a hand prosthesis. The participant attempted to perform wrist extensions with their residual limb while M-mode images, color M-mode images, and velocity estimates from each transducer were produced. This data had the unwrapping and disambiguation approach applied to mitigate aliasing as a means of demonstrating the technique on human imaging.

## IV. Results

### Simulations

#### Investigation of linear sweep parameters

As described in section II, the resolution of range and velocity is dependent upon the selection of imaging parameters. The sweep rate is determined by the bandwidth of linearly swept chirp and the sweep duration and is the most important parameter for the resolution in both range and velocity. Using the 2D Fourier Transform method to estimate the range and velocity of the simulated scatterer, the relationship between bandwidth, sweep duration, and resolution was characterized. The simulation demonstrated how with increasing sweep duration and decreasing bandwidth the resolution of the velocity improved (Fig 11 Left). Contrastingly, the resolution of the range improved with decreasing sweep duration and increasing bandwidth (Fig 11 Right). The selected bandwidth and sweep duration for imaging are also directly related to the intermediate frequency observed in the beat signal during imaging as observed in Fig 5.

**Figure 5.**
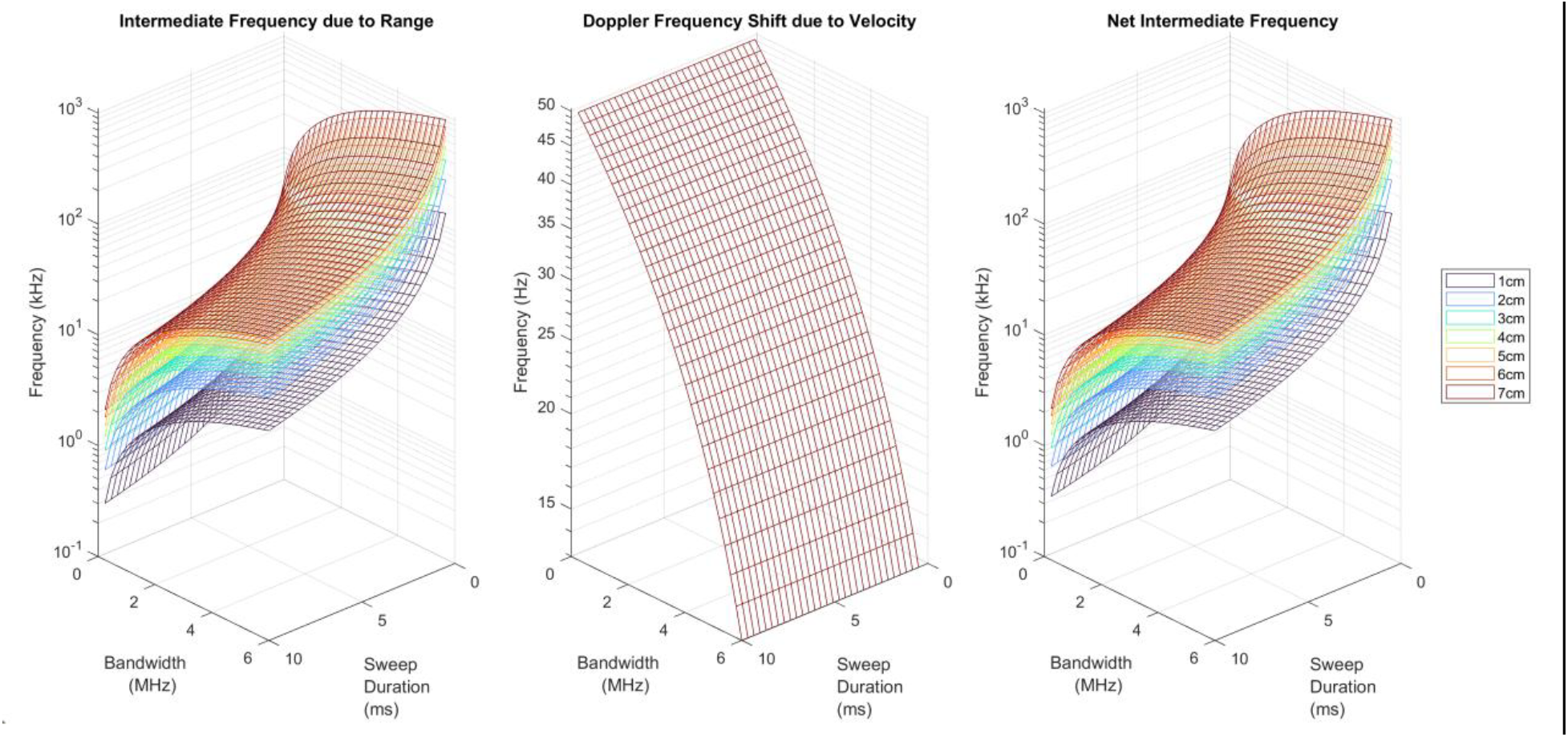
Resulting intermediate frequencies in the beat signal related to the scatterer depth, scatter velocity, bandwidth of chirps, and sweep duration of chirps. All calculations performed with a fixed scatterer velocity of 1cm/s. **Left**: Intermediate frequency due exclusively to the range of the scatter (i.e., without Doppler shift) based on sweep duration and bandwidth of chirp. **Middle**: Doppler shift of the intermediate frequency due to the velocity of the scatter. Note that the unit of the z-axis for this plot is Hz rather than kHz, and that depth of the scatterer does not impact the observed Doppler shift. Furthermore, note the log scale of the frequency axis as the Doppler frequency shift truly changes linearly with bandwidth. **Right:** Net intermediate frequency observed in the beat signal for scatterers of different ranges but with fixed velocity (1cm/s) based on bandwidth and sweep duration of transmitted chirps.

**Figure 6.**
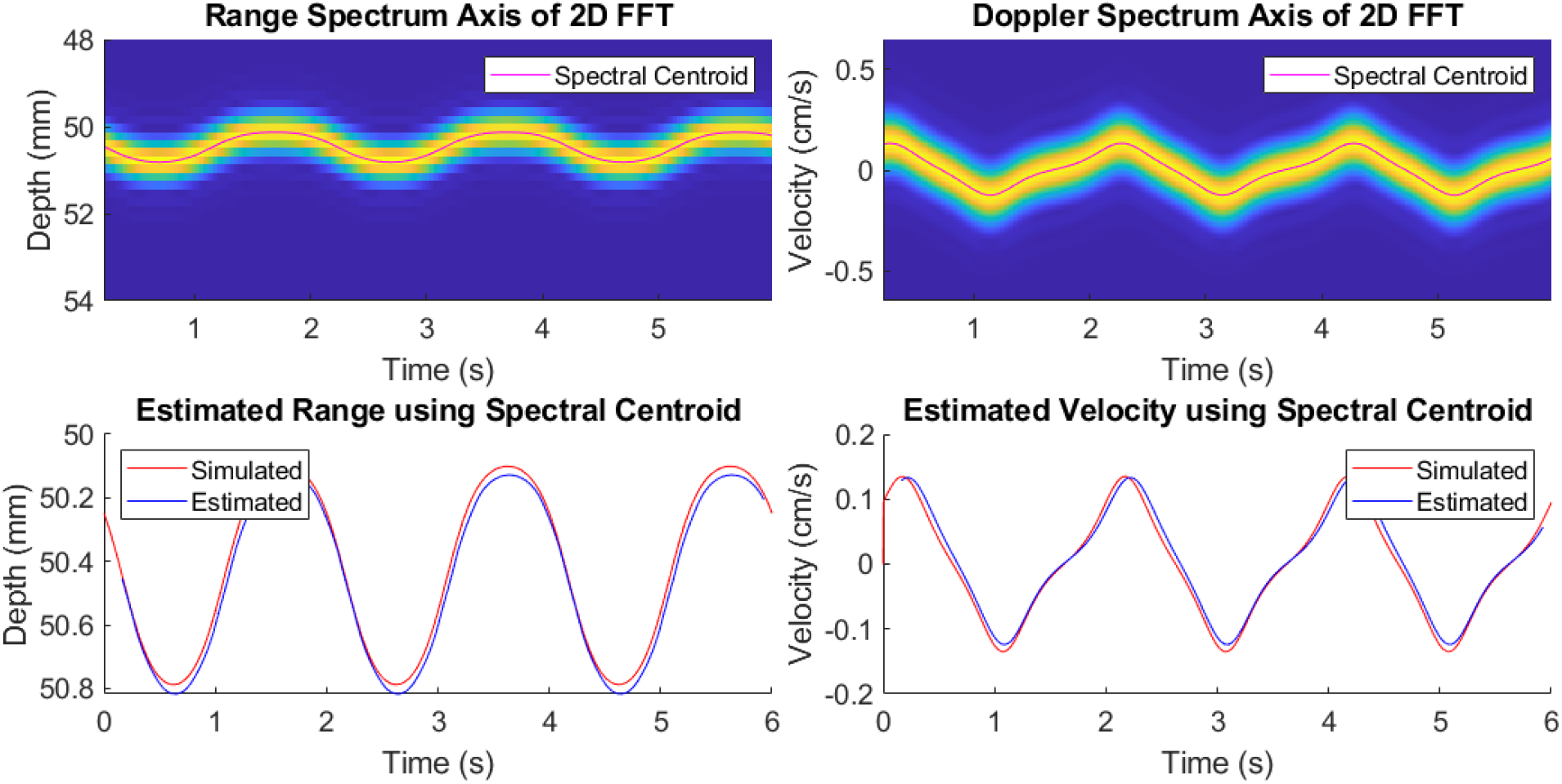
**Left Top:** The power spectral density spectrogram of the range frequency shift is reconstructed as an ultrasound M-mode image showing the scatterer position over time. The spectral centroid is overlaid on the spectrogram. **Right Top:** The corresponding power spectral density spectrogram of the Doppler frequency shift estimated from the 2D FFT is reconstructed as an ultrasound Doppler spectrogram showing the scatterer velocity over time. The Doppler frequency shifts over time follows the waveform of the simulated scatterer velocity with a processing latency corresponding to the Fourier analysis window. The spectral centroid is overlaid on the spectrogram. **Left Bottom:** The scatterer range estimated using the spectral centroid of the 2D FFT compared to the simulation’s ground truth. The estimated range waveform matches the simulated waveform with a processing latency due to the Fourier analysis window. **Right Bottom:** The scatterer velocity estimated using the spectral centroid of the 2D FFT. The estimated velocity waveform matches the waveform of the simulated scatterer velocity with a processing latency corresponding to the Fourier analysis window, however the maximum velocities are underestimated because of a finite duration analysis window of 180 ms.

**Figure 7.**
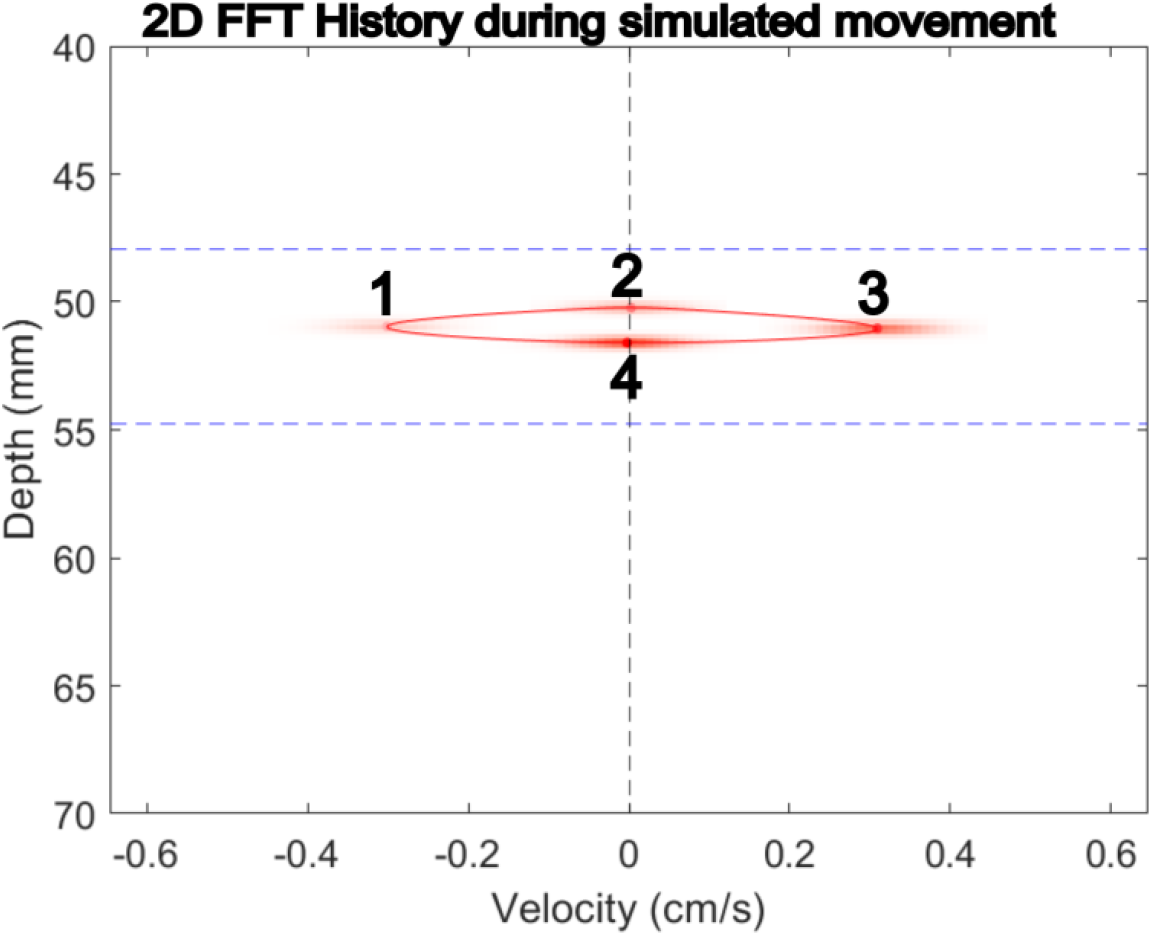
The 2D power spectral density of the same simulated scatterer from Fig 6 showing range and velocity for four time windows. The first time window (1) is presented in lower opacity than the subsequent time points, as opacity increases with each time point. Each window is constructed from ten frames of simulated imaging with a frame rate of 100Hz. The 2D spectral centroid is overlaid on the spectrograms from each time point. The black dashed line simply indicates the zero of the velocity axis, while the blue dashed lines indicate the boundaries of the depth gate used during the 2D spectral centroid calculation.

**Figure 8.**
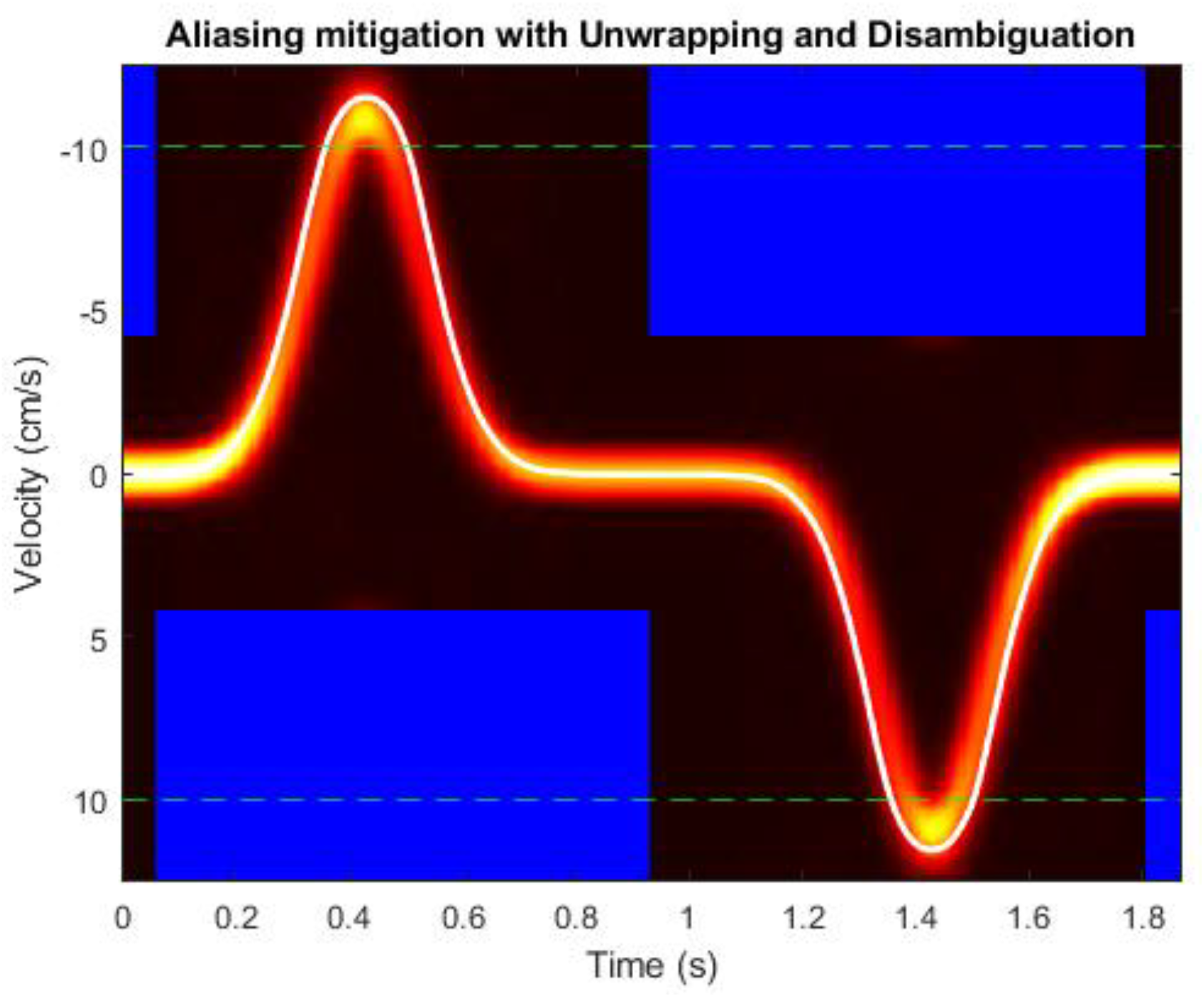
The 1D spectrogram of velocity for a simulation with a single scatterer which has had the unwrapping and disambiguation approach applied before centroid tracking. The area between the dashed green lines is the original resolvable region of the velocity spectrogram. The blue regions of the spectrogram indicate areas which have been zeroed during the disambiguation process. The white line is the final velocity centroid calculated from the unwrapped and disambiguated velocity spectrogram.

#### Investigation of the impact of noise

From the simulation investigating the impact of noise on estimating velocity as it relates to the selected sweep parameters, the RMSE for each of the velocity estimation methods were evaluated at each level of noise. The 2D Fourier transform estimation methods including the 2D spectral centroid with depth gate, 2D spectral centroid without depth gate, 1D spectral centroid, and average velocity of the color M-mode within the depth gate were broadly resistant to errors caused by noise. The observed error was mostly affected by the selection of sweep parameters. In general, the 2D spectral centroid methods had lower RMSEs with short sweep durations and narrow bandwidths with the exception being for the - 30dB SNR simulation of the 2D spectral centroid without depth gate. The 1D spectral centroid method was largely invariant to noise and sweep parameter selection at low SNRs, but at 30dB SNR and no noise sweep times of 1ms and 2ms exhibited lower RMSEs. The average velocity of the color M-mode experienced higher RMSE with lower SNRs at broader bandwidths, but the impact of this could potentially be offset by shorter sweep times.

For the autocorrelation-based velocity estimators, the RMSE for the 2D and 1D autocorrelation methods were also evaluated using different permutations of the same sweep parameters and SNRs. Both methods exhibited similar performance across all different combinations of sweep parameters and SNRs. The error was most impacted by low SNR for both 2D and 1D autocorrelation estimators particularly with short sweep durations and narrow bandwidths.

The resolution of the 2D Fourier transform was slightly impacted by noise and greatly impacted by the choice in sweep parameter. As observed in Fig 5 and Fig 9, the choice of sweep parameter greatly impacts the resolution of the 2D Fourier transform. This trend is maintained for these simulations as the longer sweep durations and narrower bandwidths resulted in finer resolution in velocity, while shorter sweep durations with broader bandwidths gave finer range resolution. When considering the impact of noise, velocity resolution gradually decreased with increasing SNR; however, at shorter sweep times the impact of noise was more significant. Similarly for range resolution, noise resulted in coarser resolution, but this effect was larger at longer sweep durations and narrower bandwidths.

**Figure 9.**
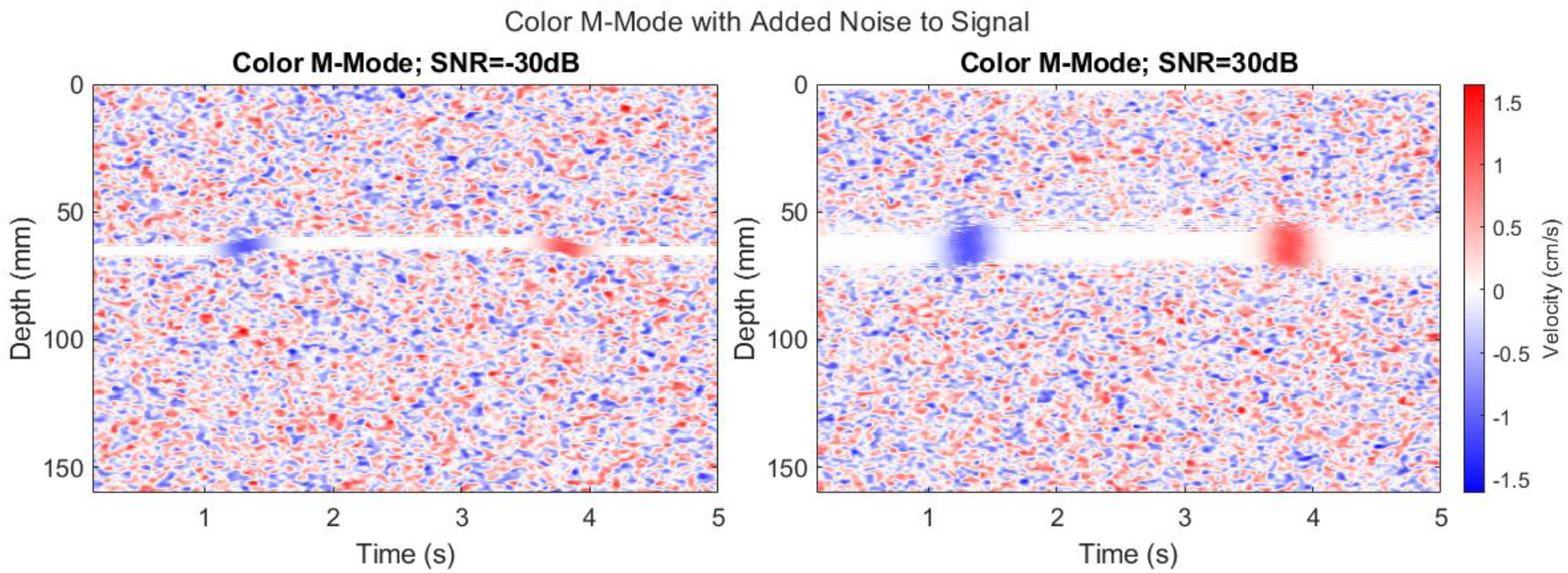
Color m-modes from a simulation of a scatterer following a trapezoidal displacement pattern with a maximum velocity of 1 cm/s at two distinct levels of noise. **Left:** Color M-mode of the simulation with an SNR of -30dB. The noise results in the random spotting of high and low velocities at all depths besides the scatterer due to changes in phase within the noise. **Right:** Color M-mode of the simulation with an SNR of 30dB. Note how the reduction in noise has resulted in broadening of the scatterer’s movement in the depth axis. This is due to spectral leakage of the Doppler shift frequency content and minimal signal from other sources at those depths stronger than this spectral leakage. When imaging tissues this spectral leakage has a minimal impact due to the presence of scatterers throughout the imaging media.

**Figure 10.**
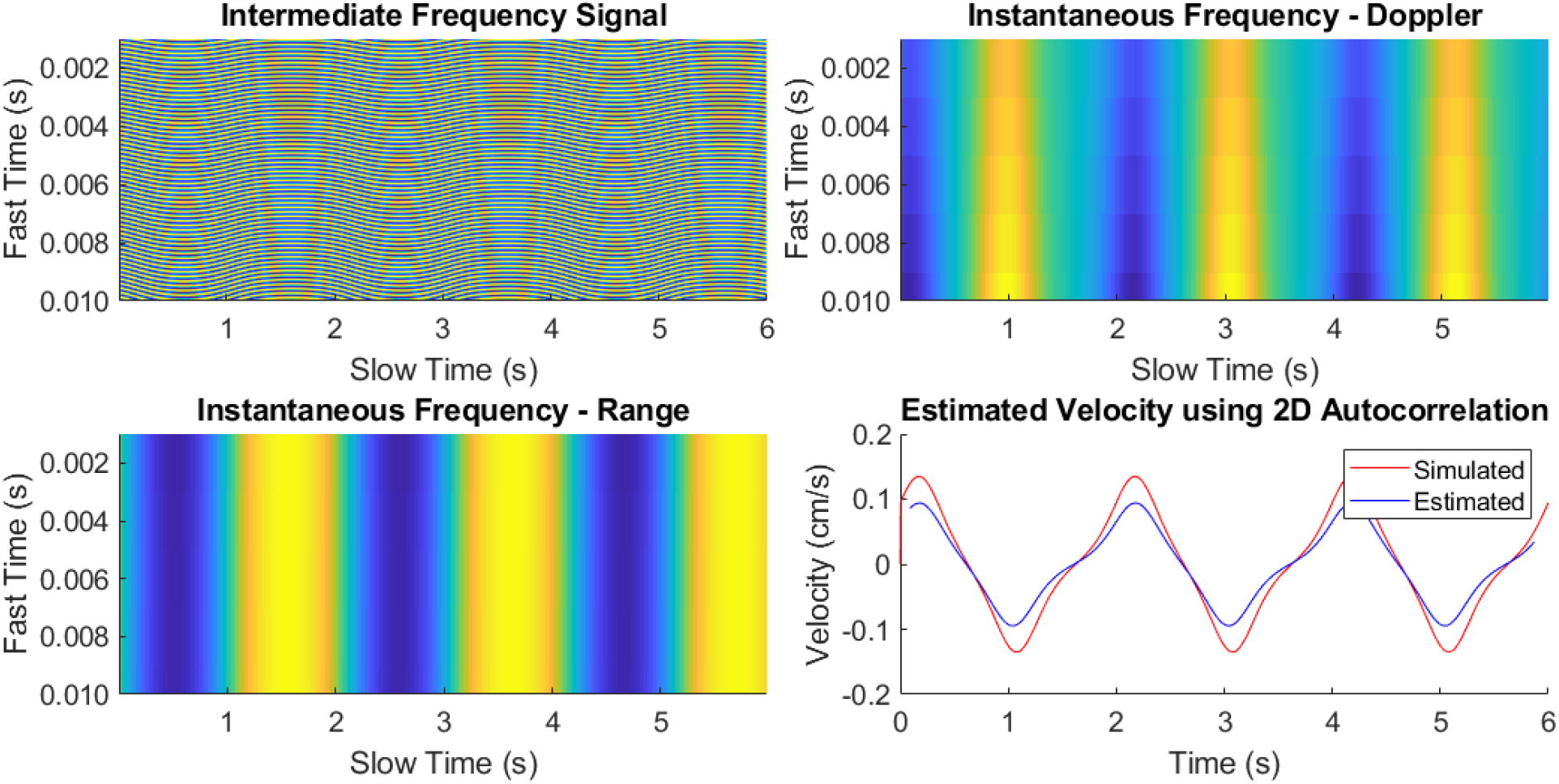
**Top Left:** The intermediate frequency signal can be represented as a 2D matrix, where the values in the rows of each column correspond to the samples of the demodulated signal within each chirp (fast time), and each column corresponds to consecutive chirps (slow time). **Top Right:** The corresponding instantaneous Doppler frequency shift estimated from the 2D autocorrelation function from Eq. 24. **Bottom Left:** The corresponding instantaneous Doppler frequency shift estimated from the 2D autocorrelation function from Eq. 25. **Bottom Right:** The scatterer velocity estimated using the lags of the 2D autocorrelation using Eq. 26. The estimated velocity waveform closely matches the waveform of the simulated scatterer velocity with a processing latency related to the autocorrelation analysis window.

**Figure 11.**
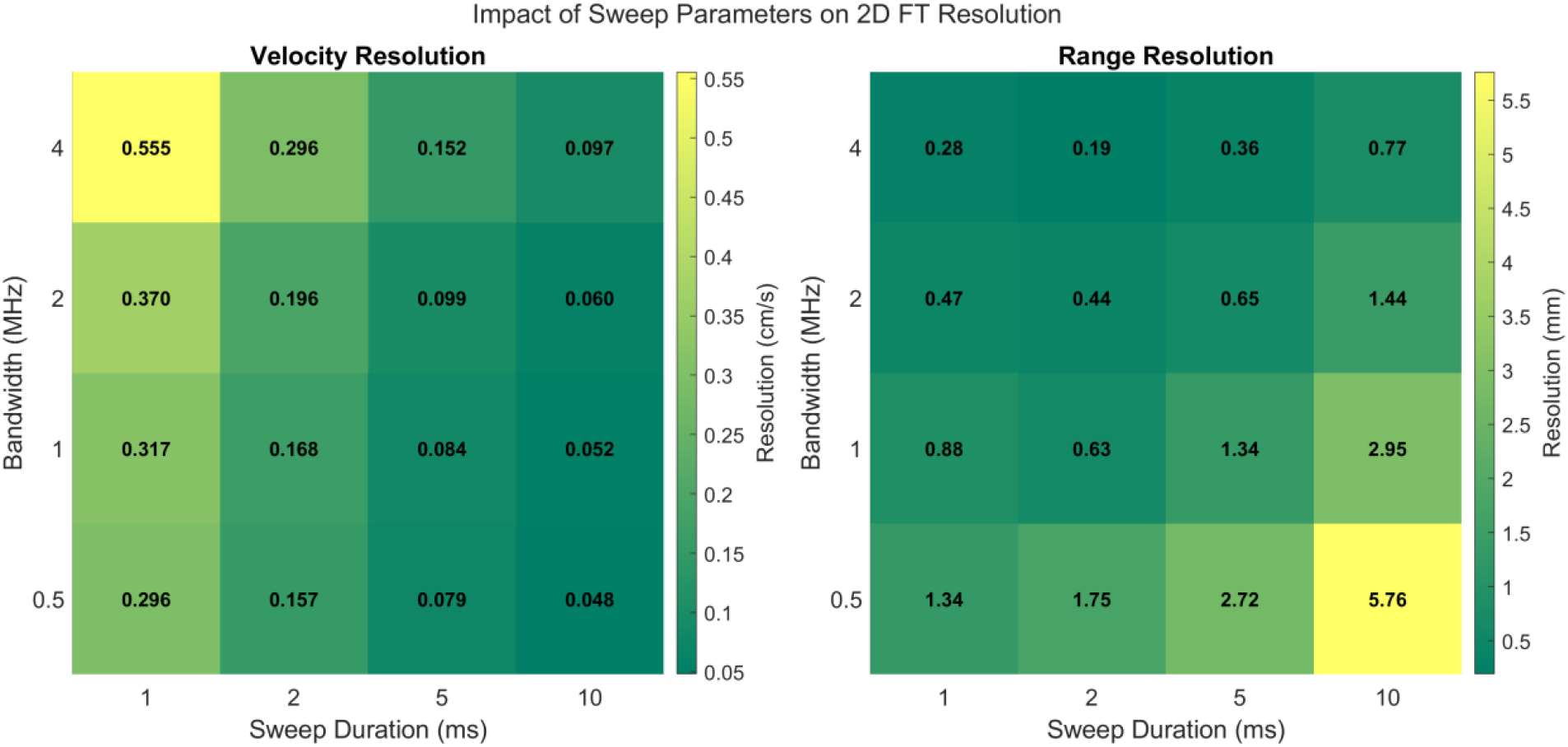
**Left:** The resolution in the velocity dimension of the 2D Fourier Transform estimation method using the spectral centroid. The decreasing bandwidth and increasing sweep duration improves resolution in the velocity dimension. **Right:** The resolution in the range dimension of the 2D Fourier Transform estimation method using the spectral centroid. The increasing bandwidth and decreasing sweep duration improves resolution in the range dimension. Increasing sweep duration and decreasing bandwidth both reduce the intermediate frequency for scatterers, simplifying necessary recording hardware, but has knock on effects for imaging quality.

**Figure 12.**
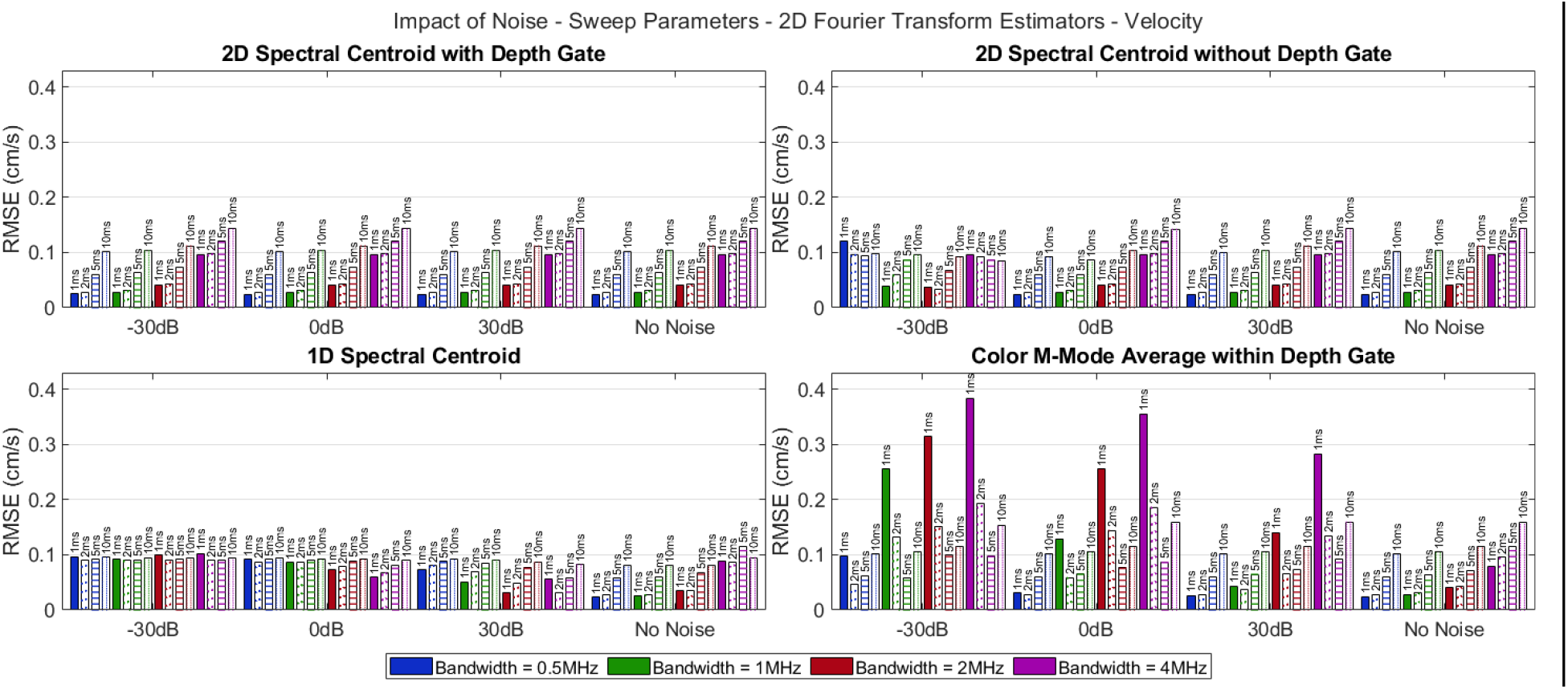
The RMSE for the velocity of the simulated scatterer using a variety of 2D Fourier transform methods including: the 2D spectral centroid with a depth gate applied, 2D spectral centroid without depth gate applied, 1D spectral centroid, and the average velocity of the color M-mode within the depth gate. Each simulation was performed with a different combination of chirp sweep duration, chirp bandwidth, and SNR.

**Figure 13.**
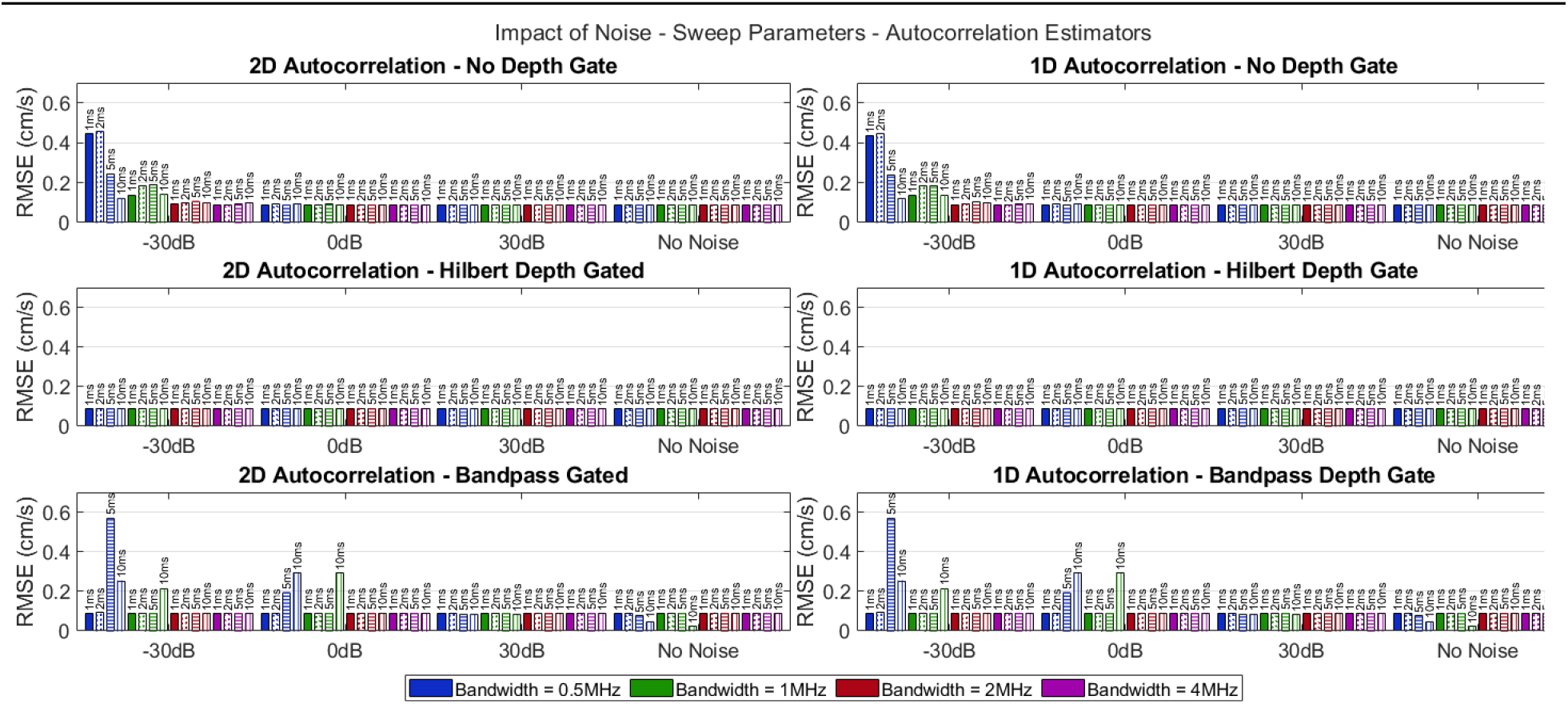
The RMSE for the velocity of the simulated scatterer using the 2D and 1D autocorrelation estimators. Each simulation was performed with a different combination of chirp sweep duration, chirp bandwidth, and SNR.

**Figure 14.**
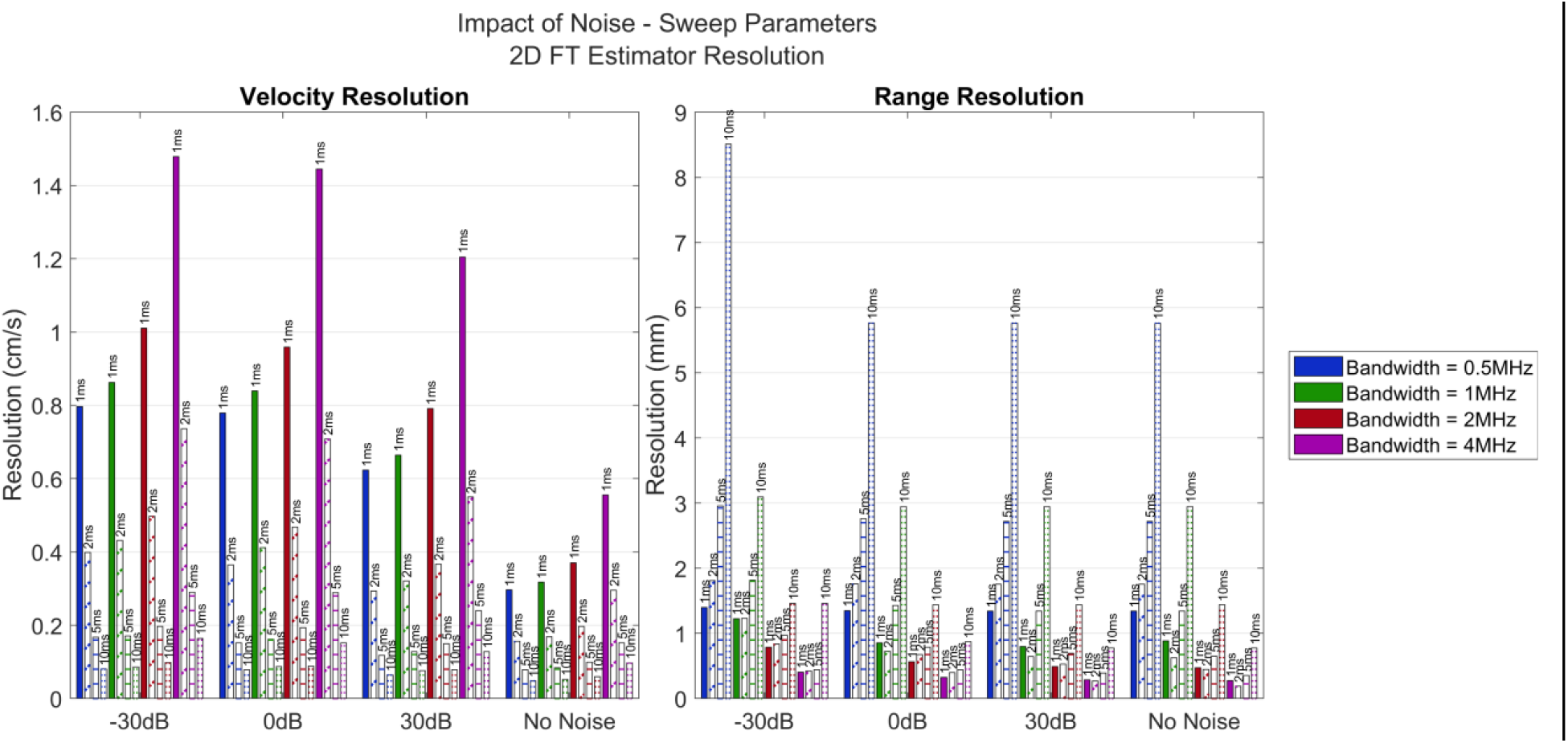
The resolution for the velocity and range of the simulated scatterer using the 2D Fourier transform. Each simulation was performed with a different combination of chirp sweep duration, chirp bandwidth, and SNR. The impact of chirp parameters has a larger impact than noise on the resolution, yet noise still results in coarser resolution.

In the simulation evaluating the impact of noise as it relates to scatterer motion, the RMSE was evaluated for both 2D Fourier transform and autocorrelation-based velocity estimation methods. When no noise was present, all the 2D Fourier transform techniques exhibited similar performance, which improved with scatters closer to the simulated transducer. With noise present the RMSE of the 1D spectral centroid increased as well as the 2D spectral centroid at -30dB. However, the other 2D Fourier transform estimators were largely resistant to noise. Moreover, the trend of lower RMSE at shallower initial depths for the scatterers was maintained.

Autocorrelation-based velocity estimators were largely resistant to noise with only marginal increases to RMSE at -30dB SNR and shallower initial depths. Across all other SNRs, RMSE was similar for each given maximum velocity. Lower maximum velocity resulted in lower RMSE for both 2D and 1D autocorrelation techniques.

The resolution of the velocity and range when using the 2D Fourier transform estimators was impacted by noise. Lower SNRs resulted in coarser velocity and range resolution than higher SNRs when scatterers had the same initial depth and maximum velocity. The velocity resolution was approximately the same across all combinations of initial depth and maximum velocity. In contrast, range resolution improved with shallower initial depths but remained relatively similar with different maximum velocities at each given depth.

### Human Imaging

#### Imaging human carotid wall movement

The imaging of the carotid wall movement produced an M-mode image using conventional TDS imaging, a color M-mode depicting velocities at all depths over time using FMSCW Doppler, and estimated velocities using the 2D Fourier Transform method. These images and the velocity are presented on the left side of Fig 15. The average velocity of the carotid arterial wall over a single cardiac cycle was calculated and presented in the right-hand side of the figure.

**Figure 15.**
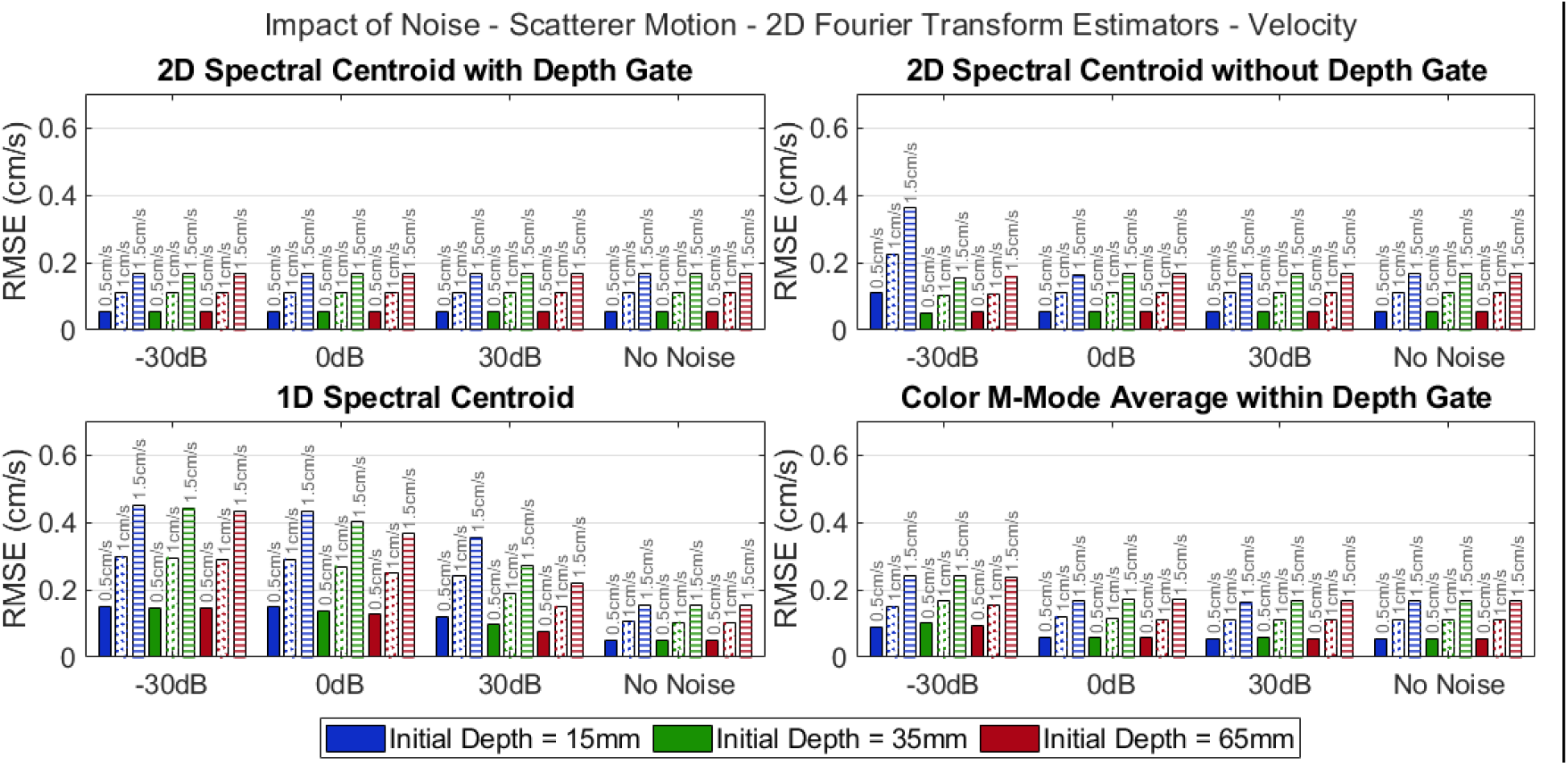
The RMSE for the estimated velocity of the simulated scatterer using the 2D Fourier transform at various initial depths and maximum velocity. Each simulation was performed with a different combination of initial depth, maximum velocity, and SNR.

**Figure 16.**
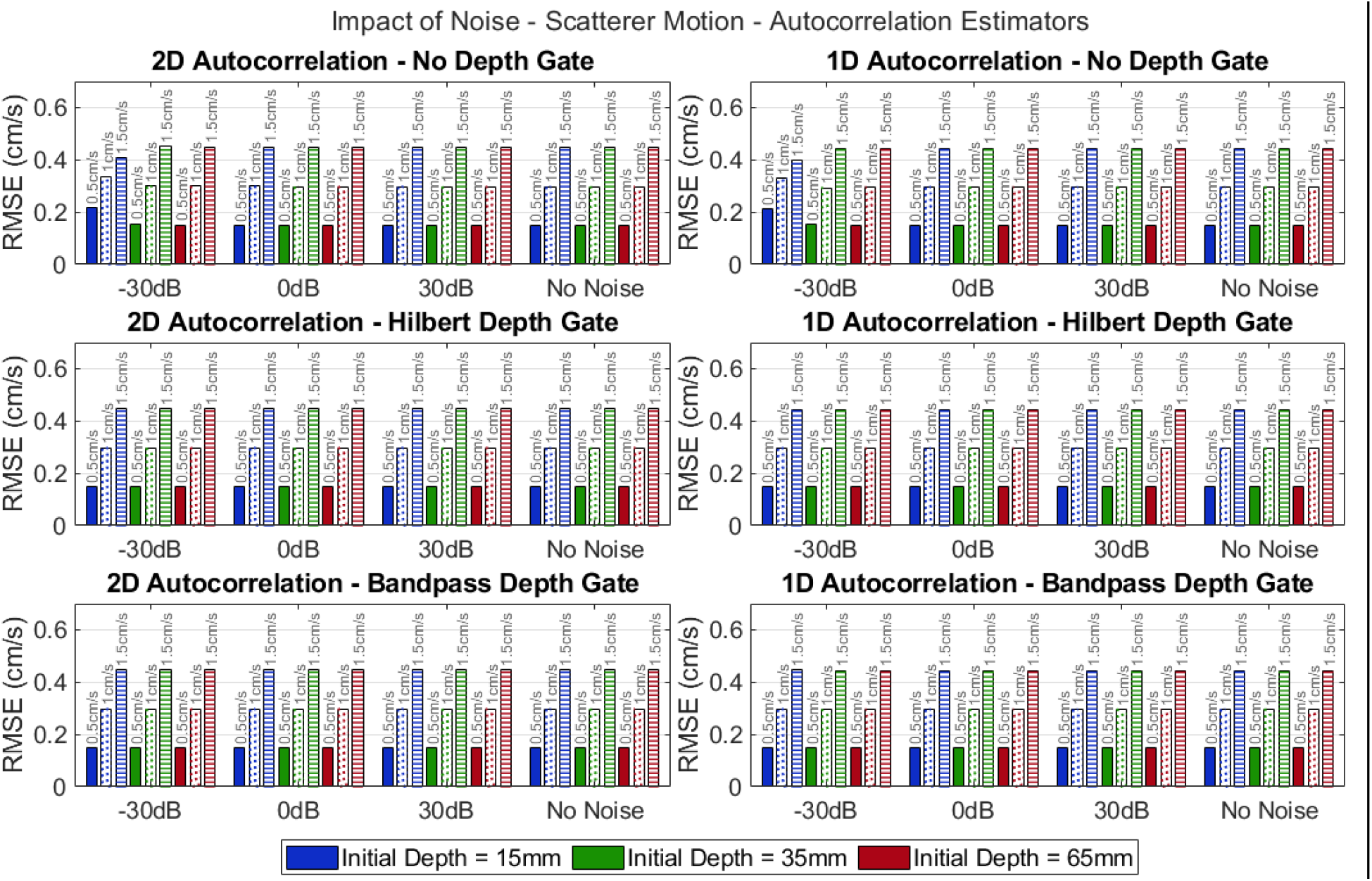
The RMSE for the estimated velocity of the simulated scatterer using the 2D and 1D autocorrelation estimators at various initial depths and maximum velocity. Each simulation was performed with a different combination of initial depth, maximum velocity, and SNR.

**Figure 17.**
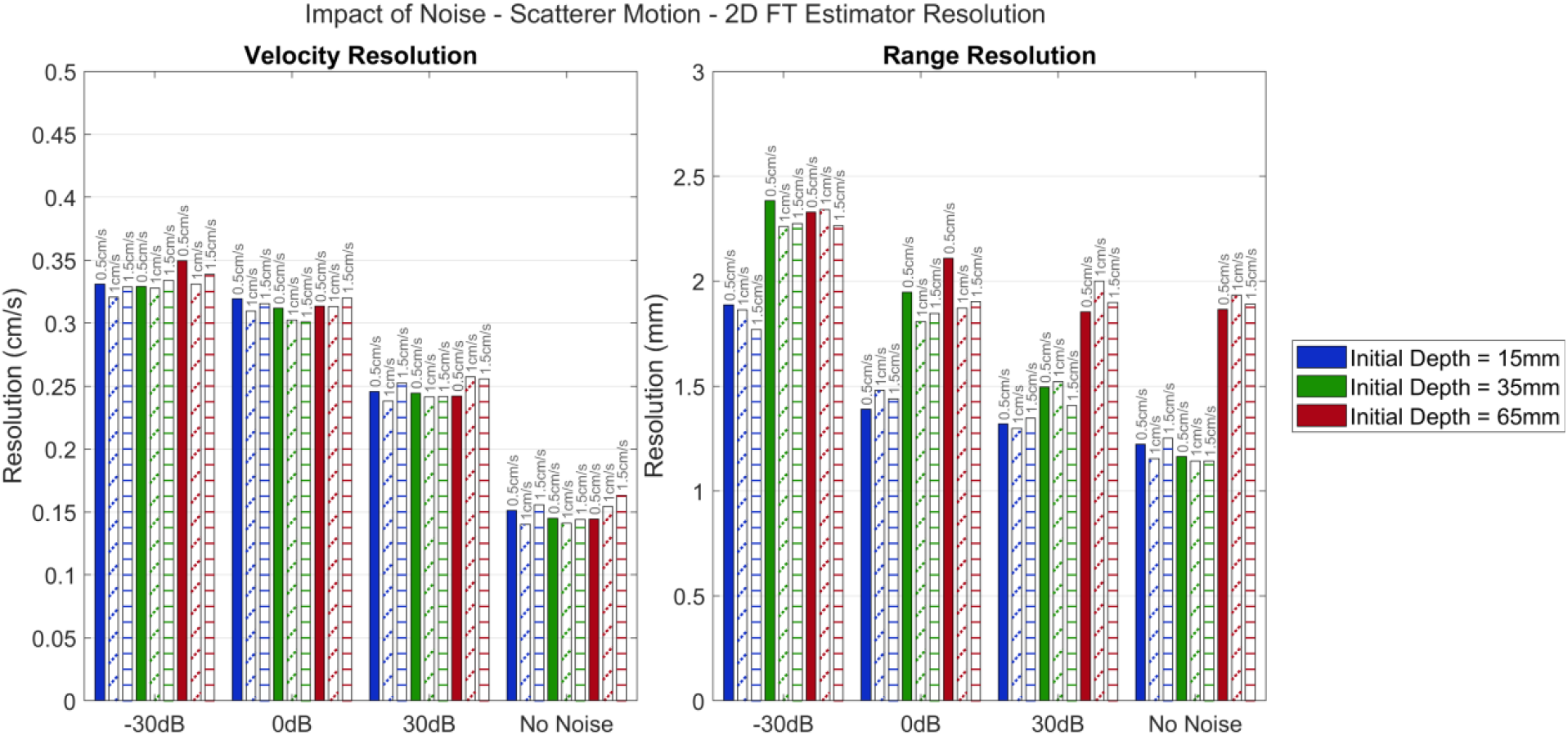
The resolution for the estimated velocity and range of the simulated scatterer using the 2D Fourier transform at various initial depths and maximum velocity. Each simulation was performed with a different combination of initial depth, maximum velocity, and SNR.

**Figure 18.**
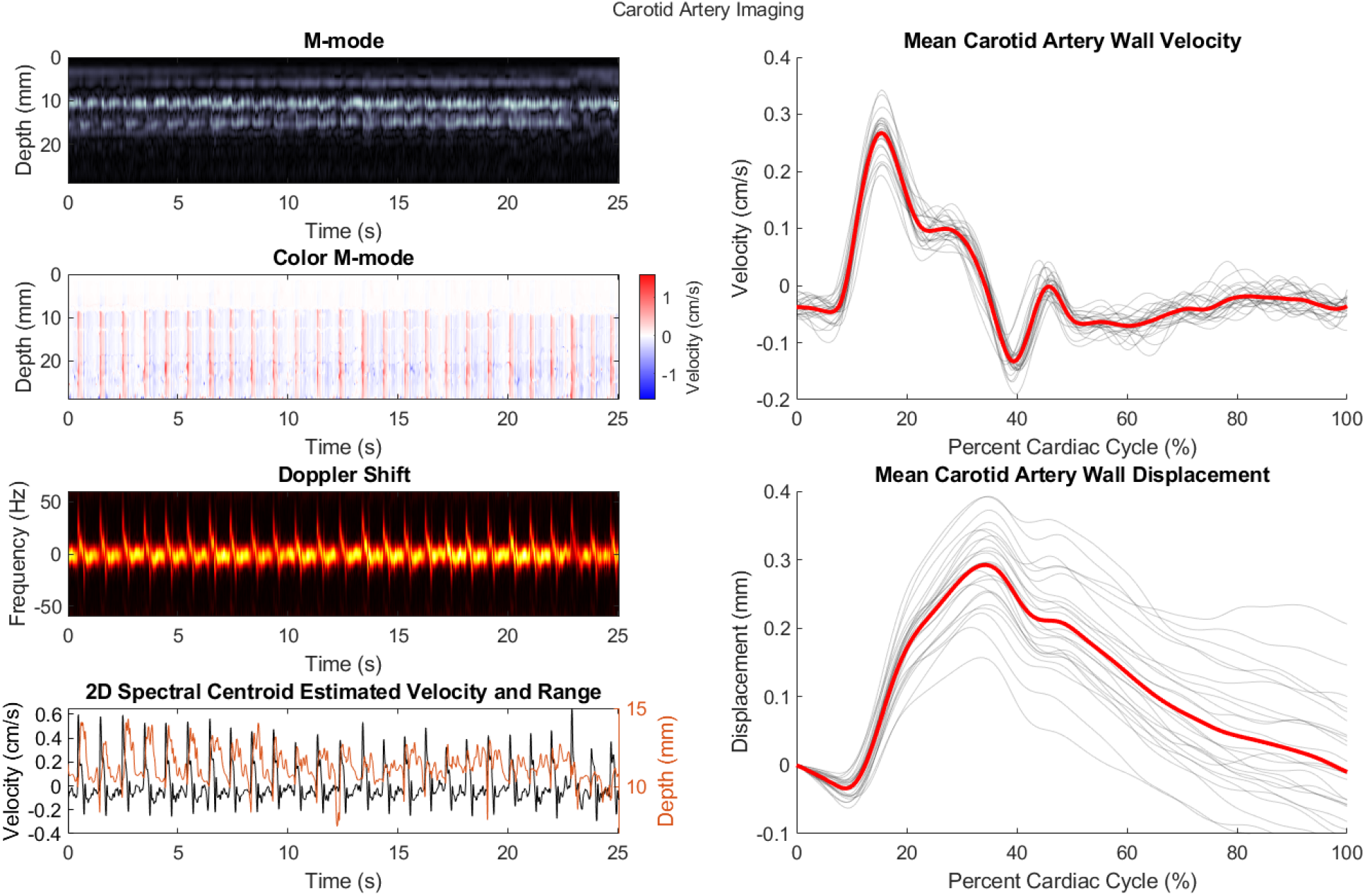
FMSCW imaging of the carotid artery wall motion over 25 seconds. **Top Left:** M-mode generated using conventional TDS imaging. **Top-Middle Left:** Color m-mode produced using FMSCW presenting the velocity at all depths being imaged. **Bottom-Middle Left:** Doppler shift due to the movement of the carotid artery. **Bottom Left:** Velocity and range (depth) estimations from the imaged carotid artery using the 2D spectral centroid. Top **Right:** Average velocity of the carotid artery during a single cardiac cycle (red) and all individual cardiac cycles which were recorded (grey). **Bottom Right:** Average displacement of the carotid artery during the cardiac cycle (red) and individual cardiac cycles which were recorded (grey). These traces were calculated by integrating the velocity traces. Furthermore, the displacement traces resemble blood pressure-like waveforms.

#### Imaging vastus lateralis movement during body weight squats

From the imaging of the vastus lateralis during body weight squats FMSCW produced a conventional M-mode image and estimates of fascia range and velocity as well as a color M-mode. The estimated velocity was utilized to produce the average peak velocity during a squat as well as the timing lag between peak contractions bilaterally and the correlation of the velocity signals at maximal alignment. The average peak velocity for the right vastus lateralis was 1.27cm/s while the left vastus lateralis had a peak velocity of 1.86cm/s. The left vastus lateralis reached its peak velocity .0125 seconds before the right vastus lateralis. The correlation between the left and right vastus lateralis velocities at maximal alignment was 0.83. The images and the velocity estimation are presented in Fig 19.

**Figure 19.**
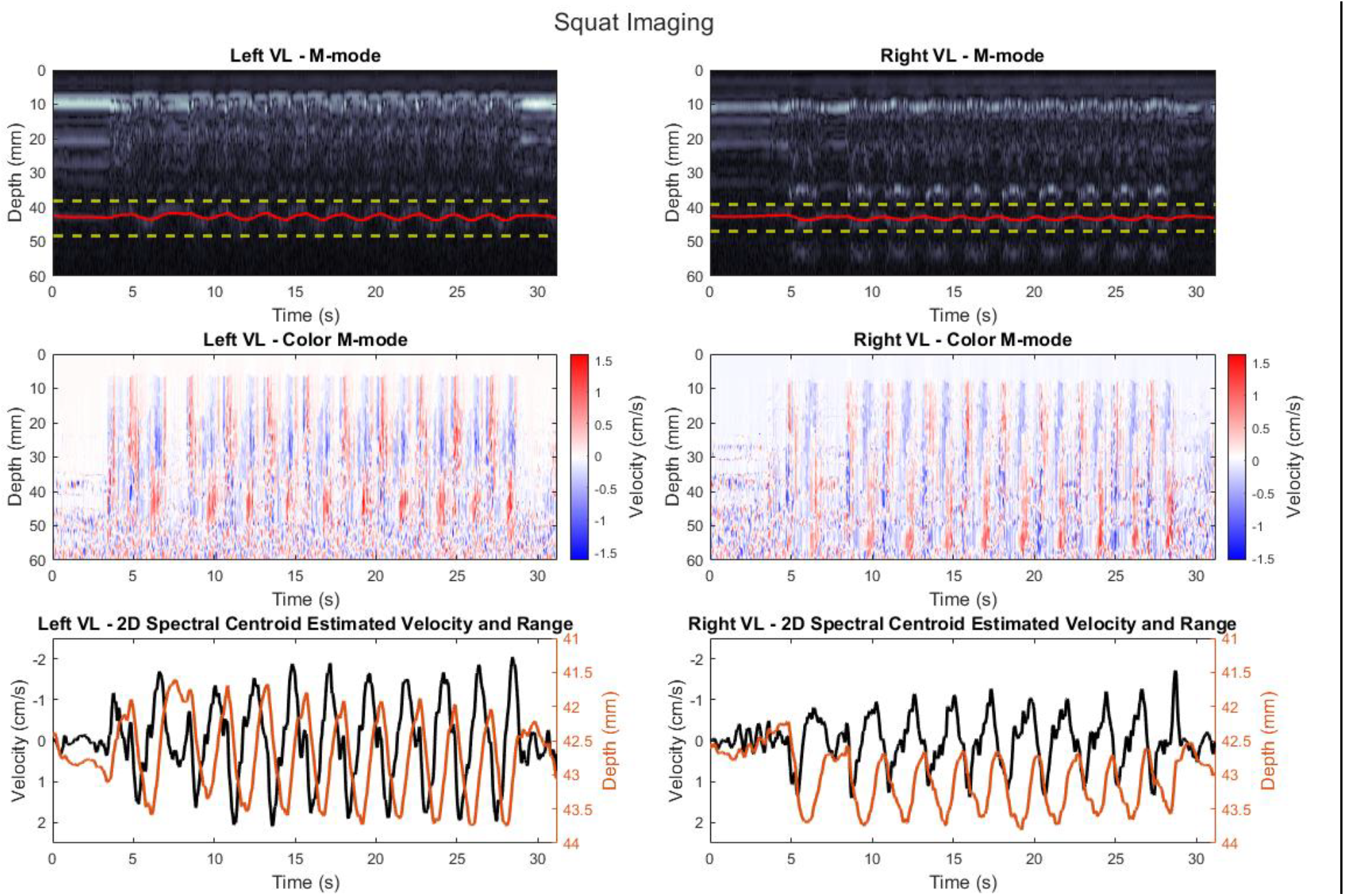
FMSCW imaging of the vastus lateralis bilaterally during 10 squats. **Top Row:** M-mode generated using conventional TDS imaging. **Middle Row:** Color m-mode produced using FMSCW presenting the velocity at all depths being imaged. **Bottom:** Velocity and range (depth) estimations from the imaged carotid artery using the 2D spectral centroid. The velocity traces (black) were utilized to produce the average peak velocity and timing lag between the two sides.

**Figure 20.**
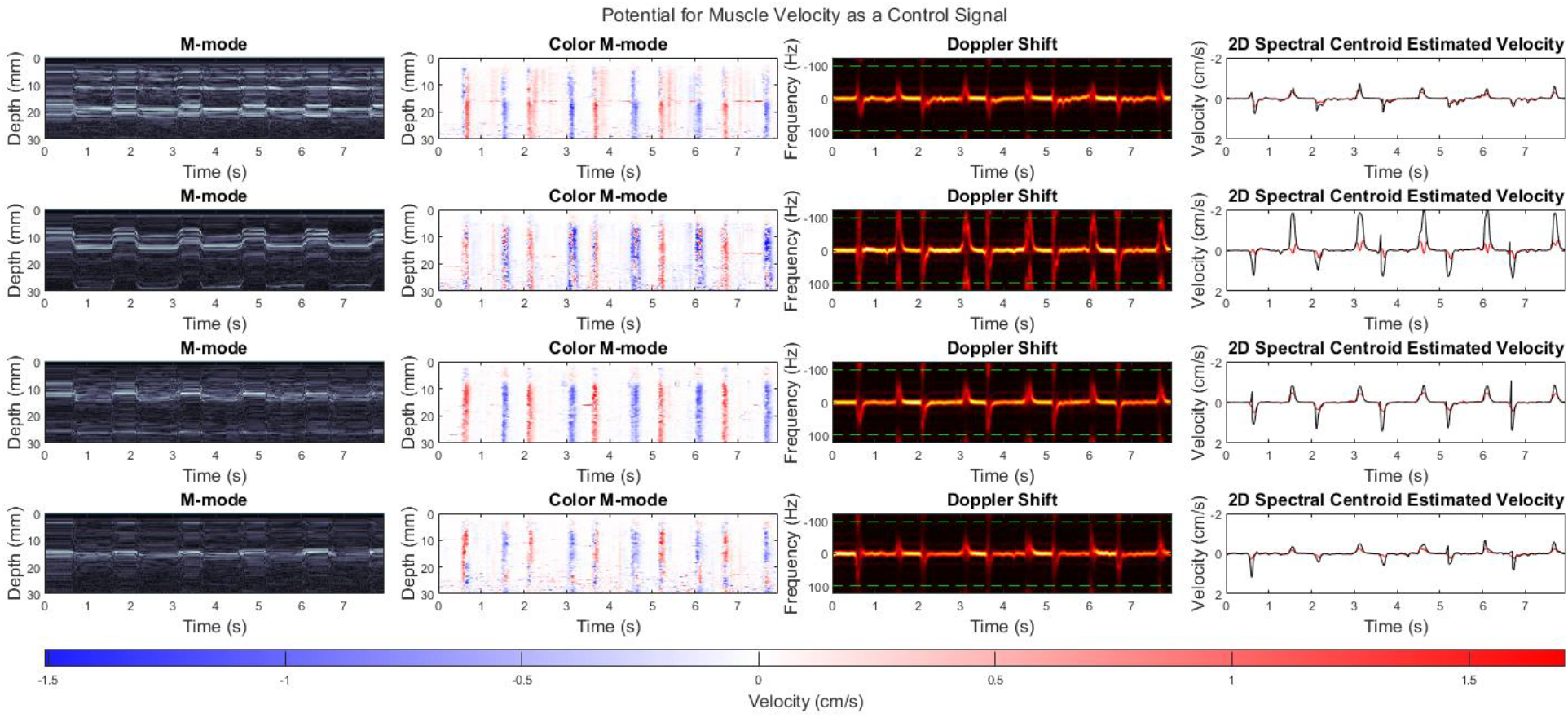
**Left Column:** M-mode images of the attempted wrist extensions of the residual limb. **Left Middle Column:** Color M-mode images of the wrist extensions. Note the clear “walls” of colors indicative of contraction and relaxation of muscle. **Right Middle Column:** Doppler shift power spectrograms clearly indicate that there is aliasing occurring in the transducer displayed in the second row as spikes in power at the top and bottom of the plot. Green dashed lines indicate the bounds of the original resolvable velocities before unwrapping. **Right Column:** Estimated velocity of each column using the 1D spectral centroid without a depth gate. The red traces are velocity centroids without the unwrapping and disambiguation approach applied to mitigate aliasing, and the black traces after this technique has been applied. The impact of aliasing can clearly be seen in the transducer displayed in the second row as there are sudden deflections in the red velocity traces in the opposite direction when the velocity exceeds that which can be resolved by the sweep parameters used for imaging. Furthermore, the color M-mode of the same transducer displays velocities of the opposite direction in each of the color walls because of the aliasing. In the black traces it can be seen how the impact of aliasing has been greatly reduced.

#### Imaging attempted hand gestures in residual limb

A third demonstration was performed imaging the residual limb of an individual with trans-radial limb difference. For this demonstration, all four transducers of the FMSCW Doppler prototype were placed on the residual limb of the participant. The participant attempted to perform wrist extension five times, and M-mode images, color M-mode images, and velocity estimates from each transducer were acquired. In this demonstration there appears to have been some aliasing in the Doppler shift which results in unnatural velocity traces. Different sweep parameters should be employed or an algorithm to adjust the mapping of frequencies to velocities based on the history of the position of the centroid should be implemented to avoid this problem in true applications.

## V. Discussion

In this study we have presented a technique for performing Doppler imaging that can be implemented using simple instrumentation that can be miniaturized into a wearable form factor. We refer to this method as FMSCW Doppler. Our proposed method has similarities to FMCW radar processing; however, our semi-continuous transmission and processing technique was designed for ultrasound applications and instrumentation. Moreover, we described how this technique exists on a continuum between traditional PW and CW methods, and by can behave similarly to PW or CW by changing imaging parameters.

Our proposed FMSCW technique is particularly well suited for wearable ultrasound applications, where data rate, hardware instrumentation complexity and power consumption are critical considerations [25]. The FMSCW approach described in this study enables an informed tradeoff between range and velocity resolution, and digitization rates as described in Figures 5 and 11. For example, the choice of the chirp sweep rate determines the range and velocity resolution (as shown in Figure 11) and also determines the range of intermediate frequencies after demodulation that determine the sampling rate needed for analog to digital conversion (as shown in Figure 5). Thus, application specific tradeoffs can be made with the same platform. Importantly, the use of long-duration swept frequencies can enable hybrid approaches between phase tracking and frequency tracking that enable the estimation of fast-moving blood flow velocities similar to frequency-based CW processing, without requiring the high pulse repetition frequencies needed by phase-based PW processing. The example demonstration applications presented in this paper utilized analog to digital sampling using 16-bit samples at 60,000 samples/s for a channel-wise data rate of 960 kbps to stream raw unprocessed data after demodulation, which can be supported by data rates of the Bluetooth Low Energy standard. Implementing the signal processing algorithms for range and velocity estimation on an embedded microprocessor can reduce the data rates significantly.

This paper presented a thorough overview of the mathematical basis of FMSCW Doppler, and the signal processing approaches used to estimate the range and velocity of scatterers. Simulations were used to evaluate the impact of selected chirp parameters and noise on resulting range and velocity estimates. FMSCW Doppler was implemented in a wearable ultrasound prototype in order to collect images from human subjects and demonstrate FMSCW in areas of potential clinical application.

The simulations performed as part of this study are aimed at evaluating the potential limits of performance for FMSCW Doppler in an ideal environment, but they also provide insight into how the structure of the technique fits into the large continuum of Doppler imaging techniques. The simulations interrogating the resolution of the 2D Fourier transform range and velocity estimator demonstrate the inherent tradeoff between the resolution of range and velocity. The selected chirp parameters determine the balance between this trade off. Velocity resolution increases with increasing sweep duration and narrower bandwidth of the chirp. This relationship is in effect indicating that sweep parameters which more closely resemble CW Doppler improve the resolution of the velocity estimate. In contrast to this, range resolution increases with decreasing sweep duration and broadening bandwidth. Selecting parameters with short sweep duration and broad bandwidth in effect pushes FMSCW Doppler to be more like conventional pulse-echo imaging techniques. However, these parameters also determine the resulting intermediate frequency produced due to scatterers. Increasing bandwidth linearly decreases the shift in frequency due to the Doppler effect (Fig 5 Middle), while the intermediate frequency due to scatterer range increases with decreasing sweep duration and increasing bandwidth (Fig 5 Left). This means that the selected sweep parameters also drive the hardware requirements to image without aliasing. Chirps with narrow bandwidth and long sweep durations result in the lowest intermediate frequency, potentially easing hardware requirements and provide finer velocity resolution; however, these sweep parameters also reduce the maximum velocity able to be estimated without aliasing. Careful consideration of sweep parameters is therefore required based on the scatterers present in the tissue of interest when applying FMSCW Doppler.

The impact of noise on the performance of FMSCW Doppler was evaluated via simulations as well. One simulation was conducted to probe the relationship between noise and the selected sweep parameters while another evaluated the interaction of noise and scatterer movement. Broadly, the sweep parameters had a larger impact on the error of the velocity estimation than the impact due to noise. This further emphasizes the importance of proper sweep parameter selection. Moreover, in the velocity estimation techniques which did see increased RMSE with noise, the sweep parameters with larger error in simulations without noise resulted in substantially larger RMSE when noise was present. 2D Fourier transform estimators using the 2D spectral centroid performed the best out of all the tested velocity estimators at all levels of noise with optimal sweep parameters. However, the RMSE of the autocorrelation-based velocity estimators, which exhibit much faster processing times, was below 0.1cm/s for most sweep parameter configurations, which may suggest that these estimators are sufficient for most real-time applications which require faster processing times.

In the simulation probing the interaction between scatterer movement and noise, it was observed that the 2D Fourier transform estimation using the 2D spectral centroid with a range gate applied was the most resistant to the impact of noise. This was the case for both range and velocity estimates. This is likely due in large part to the depth gate and the increased information from the 2D projection. The same method without a depth gate applied maintained respectable performance at most SNRs at estimating velocity but experienced greatly deteriorating performance at SNRs below 0dB for range estimates.

However, there was slight worsening of velocity estimates and substantial worsening of range estimates at low SNRs and shallow depths. This may suggest that there is an interaction between the error of the estimates and the depth of the scatterer if noise is present. The 1D spectral centroid applied to the projection of the 2D Fourier transform onto the range and velocity axes exhibited worsened performance at lower SNRs for both velocity and range estimates in comparison to the 2D spectral centroid methods; however, the 1D spectral centroid range estimate was more resistant to noise induced error than the 2D spectral centroid without a depth gate while the opposite is true for the velocity estimates. The final method based on the 2D Fourier transform was the average velocity observed in the color M-mode within the depth gate. This velocity estimation technique exhibited similar performance to the 2D spectral centroid method with depth gate at high SNRs but appeared to be unstable at low SNRs.

The autocorrelation-based velocity estimation techniques were also examined. The 2D autocorrelation estimation had a larger RMSE for all SNR values than the techniques based on the 2D Fourier transform but appeared to have relatively high resistance to noise.

Interestingly all techniques exhibited larger RMSE for scatterers at higher velocities. This is likely for three reasons: 1) the presence of the 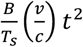 term in Eqn 12, 2) the assumption that velocity and range of scatterers is constant within a given frame, and 3) the window-based nature of the velocity estimators. The term 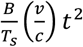 results in larger frequency spread in the intermediate frequency signal at faster velocities and ultimately distorts the resulting intermediate frequency signal making accurate estimates more difficult. Furthermore, the 2D Fourier transform estimator operates under the assumption that velocity and range of scatters is fixed within each frame. By breaking this assumption with faster velocities moving the range of scatterers or large accelerations changing both velocity and range, the frequency spread will also increase resulting in worse range and velocity estimates. The windowing function applied to the beat signal results in an underestimation of the velocity by a certain percentage of the signal amplitude as well. The impact of this effect is more extreme in the shorter slow-time (i.e., velocity) dimension ultimately producing the worsened RMSE with increased scatterer velocity.

The resolution of the 2D Fourier transform was most heavily impacted by sweep parameters rather than noise or the depth and velocity of scatterers. However, the presence of noise can dramatically worsen the resolution of the 2D Fourier transformations if suboptimal sweep parameters are utilized.

Velocity resolution was significantly coarser when noise was present with short sweep durations than with long sweep durations. Similarly, with longer sweep durations and low SNR the range resolution was much coarser than with short sweep durations with low SNR.

Demonstrations of FMSCW Doppler were performed to demonstrate the application of the technique to various domains. The imaging of the human carotid wall motion was motivated by the concept of using a worn ultrasound system for continuous monitoring of blood flow and blood pressure [26], [27], [28]. For the demonstration of our FMSCW Doppler prototype, we opted to target monitoring the movement of the carotid artery’s wall based on the imaging parameters at which our device operates. The imaging was able to estimate the velocity and depth of the wall, generating a characteristic velocity and displacement curve for the single cardiac cycle. This type of imaging may be useful for long term monitoring of blood flow or arterial health as well as research targeting blood flow in settings which would require a wearable platform such as athletics. Furthermore, while not explored as part of this study it is noteworthy that FMSCW can resolve higher velocities using a single pulse in comparison to conventional PW. As described in Eq. 30, it may be possible to estimate blood flow within a single pulse if the assumption that the vessel’s position remains fixed.

From the demonstration of the FMSCW Doppler prototype imaging the vastus lateralis bilaterally during body weight squats with a participant who had been recovering from an ACL tear, we were able to estimate the range and velocity of the primary muscle fascia within the imaging plane for each side. A depth (i.e., range) gate was used to only consider the range and velocity information within a given depth region of the tissue relative to the transducer. The resulting velocity estimations provide a wealth of information about muscle behavior during a squat including some measures related to the symmetry of the contractions, the rate of contraction, and the timing difference between the two sides. It was found that the side on which the participant had torn their ACL approximately a year before imaging was producing faster contractions than the uninjured side, but the delay was minimal and symmetry high between the two sides. While a single imaging session is unable to extrapolate many strong conclusions for rehabilitation clinicians, the results from this imaging may suggest that the injured side has been strengthened while somewhat ignoring the uninjured side. Regardless, this type of information may be utilized to evaluate the progress of rehabilitation if imaging is performed longitudinally. Monitoring the strengthening and adaptation of muscle contraction patterns poses a useful application which may specifically be suited for the application of FMSCW Doppler due to the tunable parameters to emphasize measurement of fascia depth or velocity, and the potentially simplified hardware for a wearable form factor. Beyond the estimation of the range and velocity of the primary fascia, a color M-mode may also be generated from the same data. This type of imaging provides an estimate tissue velocity from all resolvable depths. This view of muscle function can elucidate the interactions and contributions between multiple muscles within the imaging plane, which to-date has been an underexplored area of studying dynamic movements.

The imaging of the attempted movements of the residual forearm provides a notion of how a wearable Doppler system may be utilized potentially as a control signal as well as how the proposed aliasing mitigation technique works when applied to human imaging data. Sonomyographic control of prostheses is an emerging alternative to overcome limitations of conventional myoelectric control [29], [30]. However, most of these systems operate using classifiers on conventional B-mode or M-mode images. The incorporation of direct range and velocity estimations of muscle may provide a more direct control method and improve proportional control of movements based on estimated range and velocity. Proportional control schemes which use tissue Doppler may afford better fine motor control of prosthetic devices to their users. Furthermore, the introduction of a Kalman filter may further improve the performance and stability of range and velocity estimation aiding in reaching the goal of Doppler supported prosthetic control. However, the results from this imaging demonstration also highlight one of the potential pitfalls when working with Doppler imaging techniques which can alias, such as PW or FMSCW Doppler. FMSCW Doppler and PW Doppler both share the limitation of potentially aliasing if velocities exceed the resolvable Doppler frequency shift based on the selected imaging parameters.

However, the impact of aliasing may be greatly reduced if velocities are low enough such that they do not exceed 150% of the original resolvable maximum velocity in either direction. By applying the unwrapping and disambiguation alias mitigation technique we described for FMSCW Doppler in this study, we were successfully able to recover velocity traces free of the impact of aliasing from the attempted wrist movement data. In certain situations, it may be possible to ease the requirements of the pulse repetition rate if the velocities in the imaging plane are known to exist within a certain range by using the unwrapping and disambiguation approach to correct for aliasing. Yet this approach does not completely remove the impact of aliasing beyond a certain threshold of the maximum resolvable velocity. This again emphasizes the importance of the selected sweep parameters for ensuring that images and resulting velocity estimation have fine resolution and will not alias too severely.

As previously stated, FMSCW Doppler lies along the continuum of Doppler imaging techniques between the two endpoints of PW and CW Doppler, respectively. Moreover, the selection of sweep parameters drives the behavior of FMSCW Doppler towards either PW or CW Doppler. Shorter sweep times and broader bandwidths push FMSCW Doppler to behave like PW Doppler while longer sweep durations and narrower bandwidths drive it to behave like CW Doppler. Based on the needs of any given application, the sweep parameters can be adjusted such that the imaging is performed optimally to estimate the velocity or range of the scatterers. This capacity to adjust the imaging parameters to suit the needs of a given application is the greatest strength of FMSCW Doppler. Moreover, this strength of flexible resolutions can potentially be inspired by nature to adapt dynamically during imaging. Bats utilize echolocation in a manner not wholly dissimilar to FMSCW Doppler. They will use chirped transmitted pulses with long sweep durations which are able to better detect the velocity of potential prey (i.e., to ignore stable objects in the environment they echolocate for velocities in the environment which may be prey moving). Then once a potential prey target has been identified, they will change the properties of their transmitted pulses to be more akin to pulse-echo imaging with better range resolution for approaching and capturing prey [3], [18]. Biomimicry of this technique seen in bats is a potential avenue for FMSCW Doppler which presents itself as a way to adapt imaging based on the needs of the given application in real time while still utilizing one set of hardware.

## VI. Conclusion

In this study we presented the FMSCW Doppler ultrasound imaging technique in its full context to the broader field of Doppler ultrasound and how it is related to isomorphic techniques in the field radar. This Doppler imaging technique lies at a midpoint between CW and PW Doppler imaging sharing properties of both and exhibiting the ability to adjust its properties to be more like either one based on the selected imaging parameters. The key benefit of this technique over conventional PW Doppler is the simplification of the instrumentation and lower data rates since the signal processing is performed on downmixed audio-frequency signals, while the key benefit over CW Doppler is the ability to resolve range. These benefits make FMSCW Doppler particularly well suited for wearable ultrasound applications. We presented the underlying theory governing FMSCW Doppler and described and iterated upon multiple velocity estimation methods. Through simulations we demonstrated how this technique behaves with different imaging parameters, different scatter motions, and at different levels of noise. Additionally, we implemented FMSCW Doppler on a wearable ultrasound device and performed three exemplary imaging sessions to demonstrate how FMSCW may be applied to current research and clinical fields. Through both simulations and imaging of human participants, we observed and demonstrated several potential pitfalls of working with FMSCW which must be considered to achieve optimal imaging. However, the potential of FMSCW is yet to be fully realized. We believe that with inspiration from bat echolocation behavior, the technique will be able to adapt in real time to the needs of given imaging applications in order to prioritize resolution of velocity or range (i.e., depth) information.

